# LINE-1 expression in cancer correlates with DNA damage response, copy number variation, and cell cycle progression

**DOI:** 10.1101/2020.06.26.174052

**Authors:** Wilson McKerrow, Xuya Wang, Paolo Mita, Song Cao, Mark Grivainis, Li Ding, John LaCava, Jef Boeke, David Fenyö

**Author notes:** Correspondence should be addressed to (D.F.).

## Abstract

Retrotransposons are genomic DNA sequences that are capable of copying themselves to new genomic locations via RNA intermediates; LINE-1 is the only retrotransposon that remains autonomous and active in the human genome. The mobility of LINE-1 is largely repressed in somatic tissues, but LINE-1 is active in many cancers. Recent studies using LINE-1 constructs indicate that host cells activate a DNA damage response (DDR) to repair retrotransposition intermediates and resolve conflicts between LINE-1 and DNA replication. Using multi-omic data from the CPTAC project, we found correlations between LINE-1 expression and ATM-MRN-SMC DDR signalling in endometrial cancer and between LINE-1 and the ATR-CHEK1 pathway in p53 wild type breast cancer. This provides evidence that conflicts between LINE-1 and DNA replication occur in at least some human cancers. Furthermore, LINE-1 expression in these cancers is correlated with the total amount of copy number variation genome wide, indicating that, when active in cancer, pointing to a direct impact of LINE-1 associated DNA damage on genome structure. We also find that, in endometrial and ovarian cancer, LINE-1 expression is correlated with the expression of genes that drive cycle progression including E2F3, PLK1 and Aurora kinase B. This study provides evidence, supporting recent work in model cell lines, of a LINE-1/DDR connection in human tumors and raises the possibility of additional interactions between LINE-1 and the cell cycle.

## INTRODUCTION

LINE-1 (<u>L</u>ong <u>In</u>terspersed <u>E</u>lement <u>1</u>) is a family of autonomous retrotransposons that remain active in the human genome. As such, they encode proteins (ORF1p and ORF2p) necessary for the spread of LINE-1 to new genomic loci via an RNA intermediate (a phenomenon referred to as retrotransposition). ORF1p is an RNA binding protein that can form trimers and is thought to coat the LINE-1 RNA (Martin, 2006). ORF1p phosphorylation is required at several SP sites is required for retrotransposition(Cook et al., 2015). ORF2p is an enzyme with endonuclease (Feng et al., 1996) and reverse transcriptase (Mathias et al., 1991) activities that binds to the LINE-1 RNA, most likely at or near its poly A tail (Doucet et al., 2015). In dividing cells, this LINE-1 RNA/ORF1p/ORF2p ribonucleoprotein complex (RNP) can enter the nucleus during M phase and retrotranspose via a process called target primed reverse transcription (TPRT) during S phase (Mita et al., 2018). LINE-1 elements are expressed during early development (Beraldi et al., 2006; Kano et al., 2009; Percharde et al., 2018), but in somatic tissues, many host factors contribute to LINE-1 silencing, including through DNA methylation, H3K9 methylation and RNA silencing (Yang & Wang, 2016). Several studies indicate that this silencing may not always be complete, and some LINE-1 RNA can “leak through” (Belancio et al., 2010; Navarro et al., 2019), but the extent of this leak through and whether it leads to protein expression or retrotransposition remains hotly debated.

In contrast, pervasive LINE-1 de-repression occurs in ~50% of human cancers (Ardeljan et al., 2017; Rodić et al., 2014). In rare cases, de-repressed LINE-1 elements will drive cancer progression by jumping into and disrupting key tumor suppressor genes, most notably APC in colon cancer (Cajuso et al., 2019; Miki et al., 1992; Scott et al., 2016). Building on previous experiments indicating that LINE-1 can induce widespread double stranded breaks (Gasior et al., 2006) and trigger p53 mediated apoptosis (Haoudi et al., 2004), two recent papers proposed a model in which LINE-1 functionally interacts with the replication fork and factors involved in homology mediated DNA repair active during S phase, impacting the fitness of replicating cells. Mita et al. found that homologous recombination (HR) and Fanconi Anemia (FA) factors restrict retrotransposition. They also observed both that LINE-1 overexpression led to the accumulation of cells in early S phase, and that aphidicolin induced replication stress promotes retrotransposition, indicating that LINE-1 both promotes and takes advantage of replication fork stalling (Mita et al., 2020). Ardeljan et al. found that knockout of many DNA damage response genes was synthetic lethal with LINE-1 overexpression. This was observed for most of the FA group. They also observed an increase in RPA coated single stranded (ss) DNA, a marker of stalled replication forks, in response to LINE-1 expression (Ardeljan, Steranka, et al., 2020). In line with this observation Mita et al. found a functional interaction between LINE-1 activity and this ssDNA coating protein suggesting that unprotected ssDNA may be a “privileged” site of LINE-1 insertion (Mita et al., 2020).

These studies propose an exciting role for LINE-1 in cancer. Replication stress and persistent DNA damage are a double-edged sword for cancer. High levels of DNA damage are toxic to cells, with rapidly dividing cells, such as cancer cells, being the most affected. This fact explains the treatment efficacy of radiation and many chemotherapies. However, cancer cells, especially those with p53 or other DNA damage response (DDR) pathway mutations, can tolerate and even benefit from a low level of DNA damage and replication stress. Persistent DNA damage and replication stress can drive tumor evolution by promoting genome instability and copy number variation (CNV) (Gaillard et al., 2015). Therefore, if LINE-1 expression promotes replication stress and DNA damage in cancers, it would have implications both for tumor development and also for treatment. However, the Ardeljan et al. and Mita el al. studies use models in which LINE-1 is overexpressed from constructs that have been introduced to cell lines. It still remains to be shown that a similar interaction between LINE-1 expression and replication stress occurs in actual human cancers. Another recent study (Rodriguez-Martin et al., 2020), which looked at somatic LINE-1 insertions in nearly 3000 cancer genomes, did identify correlations between LINE-1 insertion and multiple classes of structural variation, including several instances where a structural variation could directly be attributed at an improperly repaired retrotransposition intermediate. These results provide support for a widespread LINE-1 / DNA damage relationship in cancer, but do not elucidate the changes in expression and signalling that are involved.

Large scale multi-omic studies (such as CPTAC) that combine DNA and RNA sequencing with shotgun proteomics and phosphoproteomics provide an opportunity to test the hypothesis that LINE-1 expression is correlated with replication conflicts and DNA damage. Dividing cells respond to DNA damage and replication stress by activating cell cycle checkpoints that pause cell division while DNA repair is ongoing. Checkpoints occur at the G1/S transition before DNA replication begins, during S phase, and at the G2/M transition before the cell commits to mitosis, and are activated through kinase signaling cascades. The integration of phosphoproteomic data allows us to measure these cascades. Furthermore, quantitative proteomic data allows us to directly measure the expression of DNA repair genes and proteins such as cyclins that mark stages of the cell cycle.

If, as the overexpression studies indicate, there is a LINE-1/replication fork conflict, higher LINE-1 expression should lead to S phase checkpoint activation. The canonical S phase checkpoint pathway is ATR-CHEK1. Activated by RPA coated ssDNA (most often occurring at stalled replication forks), ATR phosphorylates and activates CHEK1 (CHK1), activating checkpoint (Saldivar et al., 2017). However, other S phase checkpoint pathways also exist and are activated in different contexts. In the canonical response to double strand breaks, ATM phosphorylates CHEK2 (CHK2), activating checkpoint (Smith et al., 2010). Several studies indicate that ATM activation of S phase checkpoint depends on MRN and cohesin phosphorylation. First, the MRN complex (MRE11, RAD50, NBN) localizes to double strand (ds) breaks. RAD50 and NBN are then phosphorylated by ATM, a process that is necessary for ATM phosphorylation of the core cohesin subunits (SMC1A, SMC3), which in turn is necessary for activation of S phase checkpoint in response ds break inducing ionizing radiation (IR) (Magtouf Gatei et al., 2011; Luo et al., 2008; Yazdi et al., 2002). ATR activation is most associated with stalled replication forks, whereas ATM activation is most associated with ds breaks. However, the two are not independent. Homologous repair of ds breaks requires end resection, which creates ssDNA and can activate ATR. Similarly, if stalled replication forks are not quickly resolved, they can collapse, yielding ds breaks and ATM activation (Smith et al., 2010).

Because human phosphorylation sites (phosphosites) likely number in the hundreds of thousands (Vlastaridis et al., 2017), shotgun proteomics experiments can only identify and quantify a fraction of the human phosphoproteome. For any given signalling pathway, only a fraction of phosphosites are measured. Nevertheless, we were able to identify correlations between LINE-1 (ORF1p) expression and S phase checkpoint. LINE-1 expression was correlated with ATM-MRN-SMC1A pathway activation in endometrial cancer. As were several drivers of cell cycle progression, potentially indicating adaptation to chronic DNA damage. Total CNV was also correlated, indicating that this chronic stress may have serious genomic consequences. In p53 wildtype breast cancer, we see an anti-correlation between LINE-1 expression and ATM protein levels and positive correlation between LINE-1 expression and ATR-CHEK1 activation, indicating S phase checkpoint activation via a separate pathway. These correlations are absent in p53 mutant breast cancer. We do not identify correlations between LINE-1 expression and DDR related phosphorylation in ovarian cancer, but we do see a positive correlation with PARP1 protein levels and with several markers of cell cycle progression. Significant associations between LINE-1 and DDR/CNV were not observed in colon cancer, potentially for technical reasons (see discussion). We therefore give colon cancer limited consideration. In contrast to other cancers examined in this study, we did not identify strong evidence of widespread LINE-1 expression in kidney cancer.

## RESULTS

### Quantifications of LINE-1 expression, ORF1p phosphorylation, and retrotransposition are highly correlated

LINE-1 activity can be measured at several points in its life cycle, including: LINE-1 RNA expression, ORF1 protein expression, ORF1p phosphorylation and somatic insertion. We first wanted to know how these measurements relate to each other to determine the extent that one measurement can stand as a proxy for overall LINE-1 “activity.” We reanalyzed data from the CPTAC project to quantify LINE-1 mRNA, ORF1p (L1RE1) and ORF1p S18, S27, S33 and T203 phosphorylation in 5 cancer types: breast (n=94; (Krug et al., 2020)), ovarian (n=97; (McDermott et al., 2020)), colon (n=93) (Vasaikar et al., 2019), clear cell kidney (n=106) (Clark et al., 2019) and endometrial (n=88) (Dou et al., 2020). We also used the Mobile Element Locator Tool (MELT) to quantify somatic LINE-1 insertions in kidney and endometrial tumors, where matched tumor and whole blood WGS was available. WGS was not available for breast, ovarian or colon tumors. ORF2p quantifications were attempted (results not shown), but did not correlate with ORF1p, consistent with our previous observation that endogenous ORF2p is not readily detectable by shotgun proteomics (Ardeljan, Wang, et al., 2020). Because LINE-1 is subject to regulation throughout its life-cycle and different quantifications measure different life-cycle stages, they may be uncorrelated or weakly correlated in certain contexts. Figure 1 shows how these quantifications measure different points of the LINE-1 life-cycle: LINE-1 mRNA may be present as ‘stand-alone’ (B) transcripts or as RNPs co-assembled with ORF1p and ORF2p (C,D); ORF1p, which may be present as part of an LINE-1 RNP or separate from the LINE-1 RNA (not shown) is primarily cytoplasmic (C), but can also be found in the nucleus (D) in G1 phase (prior to retrotransposition which occurs in S phase) (Mita et al., 2018); somatic insertions indicate completed cycles of retrotransposition (E).

**Figure 1.**
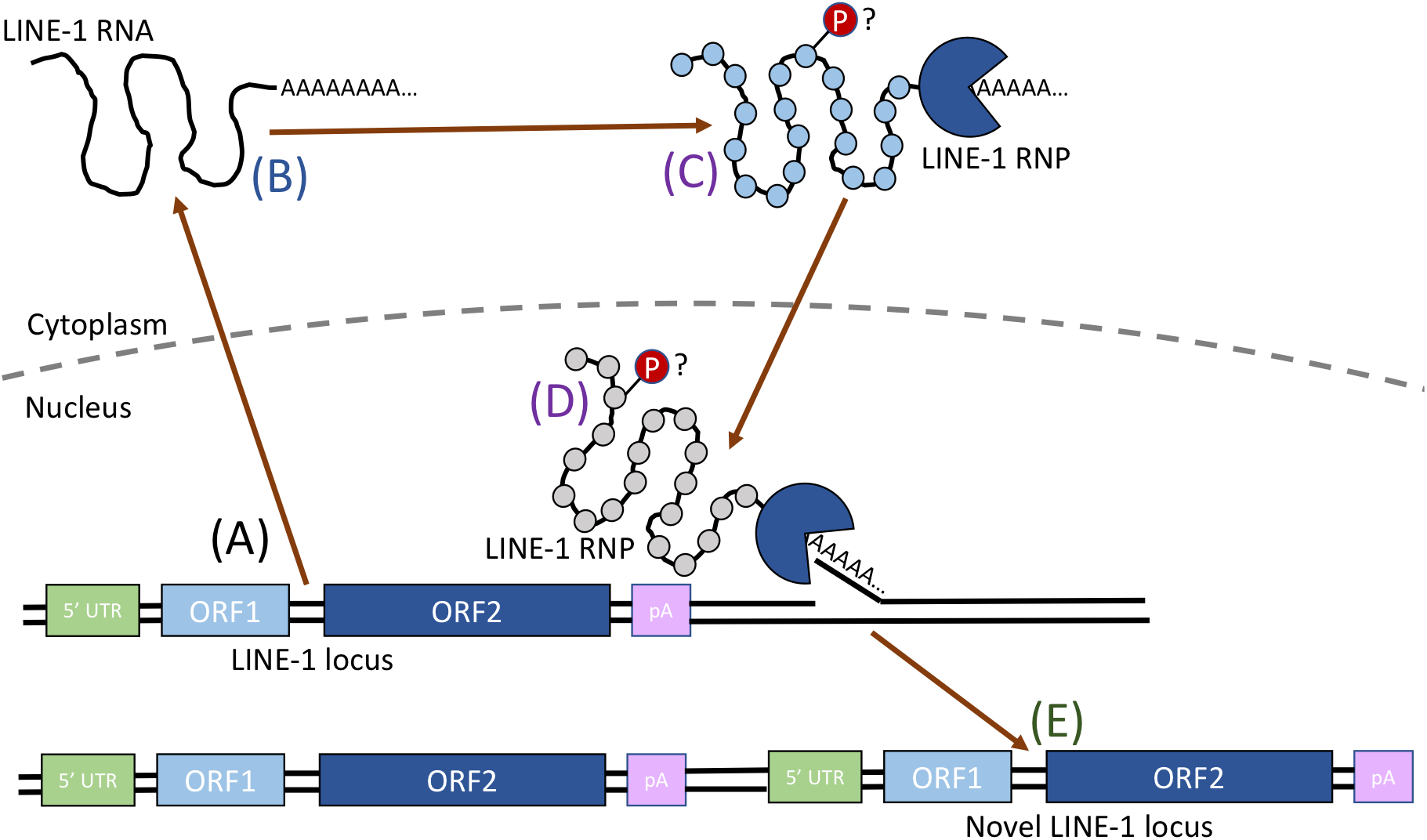
Schematic showing the LINE-1 lifecycle and the points at which LINE-1 products are quantified. (A-E) shows the path of retrotransposition. LINE-1 RNA may be loose (B), in a cytoplasmic RNP (C) or in a nuclear RNP (D). LINE-1 ORF1p may be in a LINE-1 RNP (C,D), or isolated (not shown). ORF1p in the nuclear RNP (D) is shown in gray as it is present when the LINE-1 RNP enters the nucleus, but is removed before S phase. The timing of ORF1p phosphorylation is unknown, but likely occurs at the LINE-1 RNP stage (C,D) as it seems to play a role in retrotransposition. Novel insertions (E) indicate completed LINE-1 retrotransposition events.

Overall, we find strong correlation among all four measures of LINE-1 activity: LINE-1 RNA, ORF1p, ORF1p phosphorylation and somatic insertion (endometrial cancer only). LINE-1 RNA and ORF1 protein correlate very well with each other (Figure 2A-E) except in kidney cancer (Figure 2D), which we attribute to low LINE-1 expression. Indeed, no more than one high confidence somatic insertion was identified in any kidney sample analyzed here. A similar lack of retrotransposition was observed in TCGA sequenced clear cell kidney tumors (Helman et al., 2014). Given this lack of LINE-1, we excluded kidney cancer from most of the subsequent analyses. For the other four cancer types, we found that LINE-1 RNA/protein Spearman correlations (ρ values) range from 0.55 to 0.79. Despite the challenges associated with quantifying the expression of repetitive elements, LINE-1 RNA/protein correlations far exceeded median RNA/protein correlations, generally around ρ=0.45 (Figure S1). We also found a significant correlation between LINE-1 protein expression and somatic insertions in endometrial cancer (ρ=0.38, p=0.004; Figure 2F), the only one of these cancer types for which matched WGS was available.

**Figure 2.**
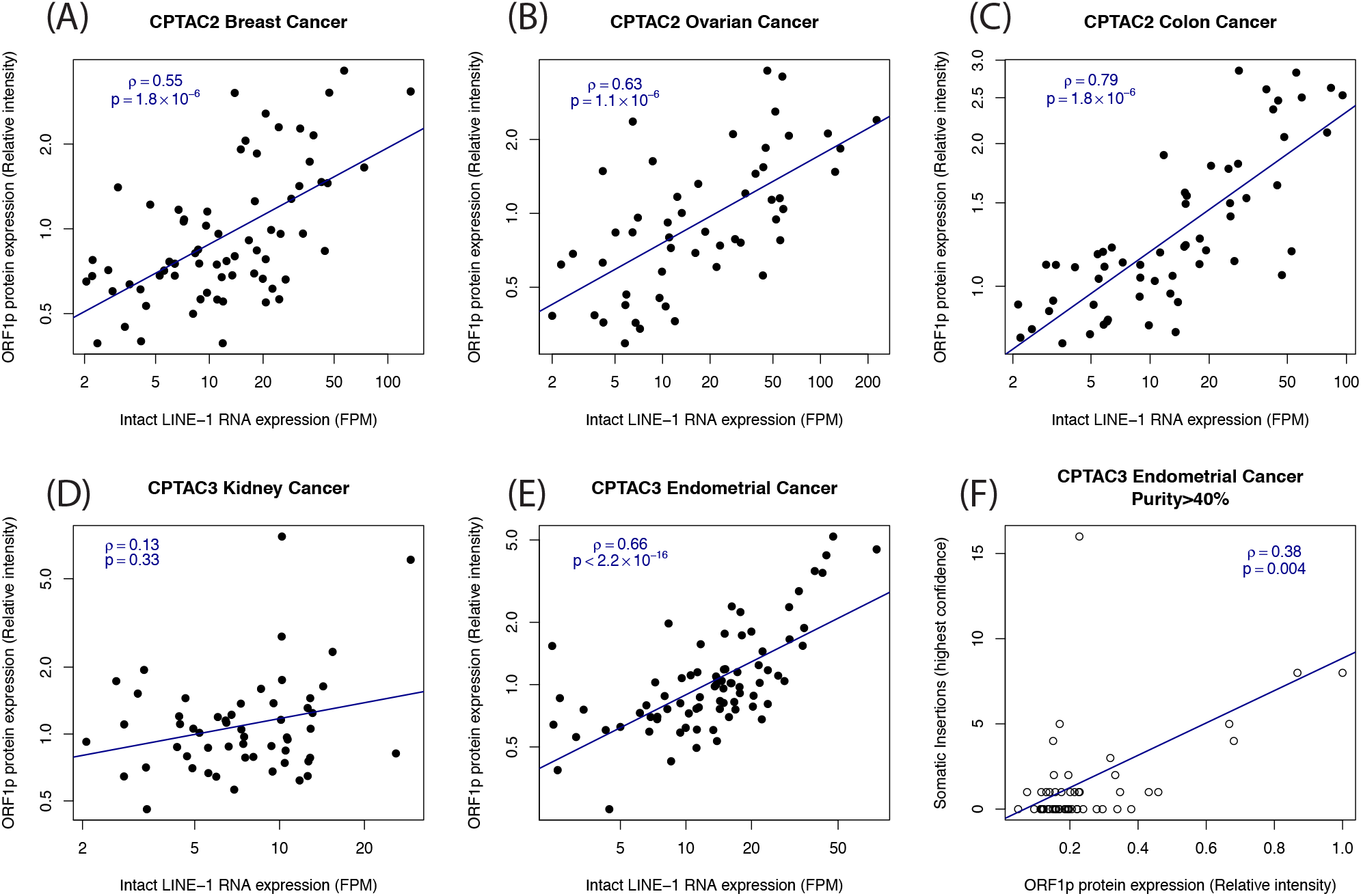
Correlation between LINE-1 RNA, ORF1p and insertion. (A-E) show LINE-1 RNA/ORF1p correlation in breast, ovarian, colon, kidney and endometrial cancers respectively. (F) Shows correlation between LINE-1 ORF1p and high confidence somatic insertions in endometrial cancer. These high-confidence insertions may be only a fraction of the total somatic insertions.

We consistently identified phosphorylation at four sites in ORF1p: S18, S27, S33 and T203 . Three of these (S18, S27 and T203) were identified in a study finding that mutation at these sites inhibits retrotransposition (Cook et al., 2015). S18 and S27 were also identified in IP-MS from colorectal cancer (Ardeljan, Wang, et al., 2020). The fourth site, S33, has not been described previously. Except in two cases where a particular phosphosite is detected never or rarely in a particular cancer type (T203 in colon cancer and S33 in endometrial cancer), these phosphosites correlate strongly with ORF1p abundance, with Spearman correlations mostly falling in the 0.5 to 0.7 range (Figure S2). The previously identified sites S18, S27, and T203 are all [S/T]P, the target for proline dependent protein kinases (PDPK). The strong correlations with substrate abundance made it impossible to use correlation to identify potential PDPKs that might target these sites, but our results nevertheless represent the strongest evidence to date that ORF1p is phosphorylated in vivo.

Overall the agreement between these LINE-1 quantifications indicates that we are able to accurately measure LINE-1 in this dataset. We therefore put most of our focus on the LINE-1 ORF1p quantification as a measure of LINE-1 expression. Further, the particularly high correlation between LINE-1 RNA and ORF1p supports the hypothesis that ORF1p binds in cis and protects its own RNA. If most LINE-1 RNA and ORF1p are part of LINE-1 RNPs, it would lead to a tight stoichiometry between the two.

### Analysis of LINE-1 RNA shows highest expression in ovarian cancer and high expression of the “hot” LINE-1 located at 22q12.1 in the TTC28 gene

Our ORF1p quantifications are relative to an internal standard that varies between cancer types and prevents straight-forward comparison of different proteins in the same sample. Thus, it is not possible to directly compare ORF1p expression across cancers. However, it is still possible to compare RNA quantifications because read counts can be compared across genes and samples if properly normalized. We find that LINE-1 RNA is most abundant in ovarian cancer, and least abundant in clear cell kidney cancer (figure 3A). Some caution should be exercised as the RNA sequencing for endometrial and kidney cancer was done using shorter reads, but at greater depth. In the case of endometrial cancer, RNA from adjacent normal tissue was also available. We find intact LINE-1 RNA to be much less abundant in normal tissue than in any of the five cancer types, including kidney (figure 3A).

**Figure 3:**
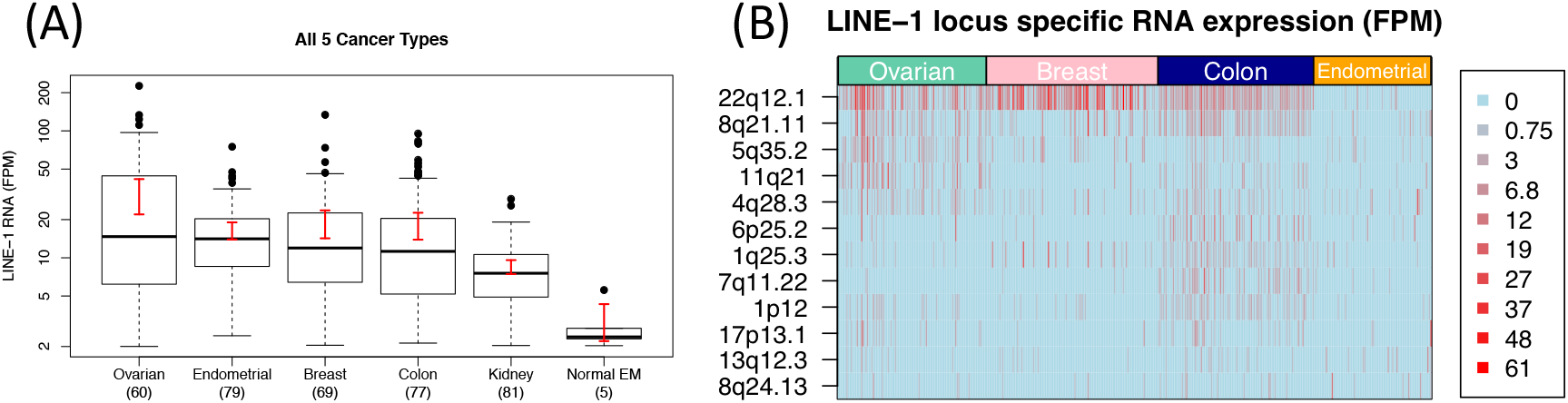
Insights from analysis of LINE-1 RNA. (A) Comparison of LINE-1 RNA quantification across cancer types. Expression is highest in ovarian cancer. (B) Highly expressed (“hot”) intact LINE-1 loci. Rows indicate an intact locus that is highly expressed (top 5) in at least one cancer type. Columns are individual cases. Darker shading indicates greater expression. Across samples/cancers, a locus at 22q12.1 is most highly expressed, accounting for two thirds of all intact LINE-1 expression in breast cancer.

Because the tool we used to quantify LINE-1 RNA, L1EM, assigns RNA to specific loci, we were able to compare locus specific LINE-1 RNA expression between cancer types (figure 3B). However, it should first be noted that L1EM assigns all LINE-1 expression to reference loci despite the presence of potentially active, polymorphic, non-reference LINE-1 loci. Thus it must be kept in mind that some LINE-1 RNA reads assigned to a locus, x, may actually be derived from a non-reference locus (or loci) for which x is likely the parent. In breast, ovarian and colon, the most highly expressed intact (full length, lacking nonsense mutations in ORF1 and ORF2) LINE-1 locus in the hg38 reference genome is located at 22q12.1, antisense to an intron of the TTC28 gene. This locus has been previously pinpointed as a highly active (“hot”) LINE-1 in several cancer types (Jung et al., 2018; Tubio et al., 2014). Among significantly expressed loci (≥2 read pairs per million), this locus accounts for 27% of intact LINE-1 expression in ovarian cancer samples, 67% in breast cancer samples and 30% in colon cancer. However, the 22q12.1 locus only accounts for about 1% of intact LINE-1 expression in endometrial cancer, where no one locus consistently dominates LINE-1 expression. A locus at 4q28.3 is the most highly expressed, yet it still only accounts for 4.5% of intact LINE-1 expression across the endometrial cancer samples.

### In endometrial cancer, LINE-1 expression is associated with p53 mutation, ATM-MRN-SMC signalling, CNV and drivers of cell cycle progression

We next turned our focus to identifying correlations between LINE-1 expression and host factors, with particular interest in whether LINE-1 expression is associated with S phase checkpoint. We begin with an analysis of endometrial cancer. To get an overview of LINE-1 expression we grouped our LINE-1 ORF1p expression estimates according to the molecular subtypes proposed by TCGA (Cancer Genome Atlas Research Network et al., 2013). This analysis revealed that most of the high LINE-1 expressing tumors fall into the “CNV high” classification (Wilcox p=6.4×10^−6^, figure 4A). As the name suggests, CNV high tumors have a high degree of copy number variation (CNV) genome-wide and have frequent p53 mutations (Cancer Genome Atlas Research Network et al., 2013; Dou et al., 2020). This is consistent with previous observations that LINE-1 expression is higher in p53 mutant tumors (Rodić et al., 2014; Wylie et al., 2016), an observation that suggests an association between LINE-1 and persistent DNA damage. Indeed, in this study we found that ORF1p levels were twice as high in p53 mutant endometrial cancers (p=0.0014). Reflecting enrichment in CNV high subtype, our LINE-1 ORF1p quantification (hereafter called LINE-1 expression) is highly correlated with genome-wide CNV (Spearman ρ=0.44, p=3.6×10^−5^, figure 4B). Strong correlation between the somatic LINE-1 insertions and structural variation was recently reported in a pan cancer study of whole genomes (Rodriguez-Martin et al., 2020). In our data, this correlation is only partially explained by p53 mutation (partial Spearman ρ=0.31, p=0.004), suggesting that LINE-1 may have a more direct role in CNV generation and that this effect may be enhanced by the effect of p53 mutation/deletion.

**Figure 4:**
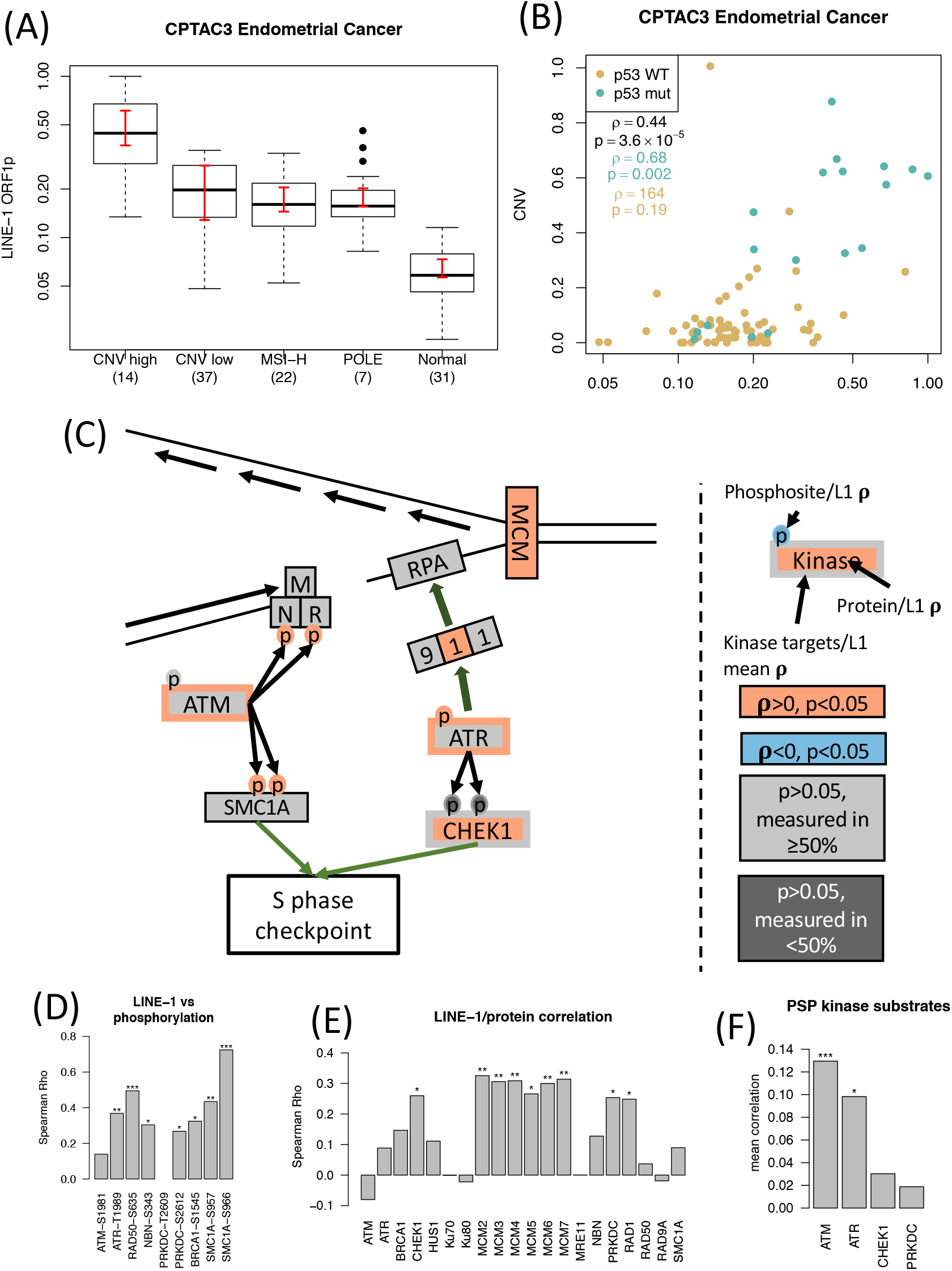
LINE-1/DDR correlations in endometrial cancer. (A) ORF1p quantification by molecular subtype: CNV high, MHV low, microsatellite instability (MSI) high, and DNA polymerase ε. LINE-1 expression is highest in CNV high endometrial cancers. (B) ORF1p vs global CNV. High CNV, high LINE-1 and p53 mutation tend to appear together. (C) A candidate model to explain the observed LINE-1/DDR correlations. At the top is a broken replication fork, on the left is the ATM-MRN-SMC pathway and on the right is the ATR-CHEK1 pathway. A protein or phosphosite filled in red indicates positive correlation at p<0.05, gray indicates no significant correlation, with dark gray indicating at least 50% missing values. Kinases are outlined in red if there is significant enrichment for correlation between ORF1p and kinase targets, and gray if not. Big green arrows indicate recruitment; small green arrows indicate downstream effects. (D) Spearman correlation between ORF1p and phosphosites pictured in C. *:0.01<p<0.05, **:0.001<p<0.01, ***:p<0.001. (E) As in D, but for ORF1p/protein correlation. (F) KSEA enrichment for correlation between ORF1p and kinase targets.

We next examined the phosphoproteomic data for correlations with S phase checkpoint. We calculated Spearman correlation between our ORF1p quantifications and host protein phosphosites identified and quantified in the CPTAC phosphoproteomic analyses (table S3). Among phosphosites with a known downstream effect annotated in the phosphositeplus database (Hornbeck et al., 2015), RAD50-S635 (ρ=0.49, p=5.6×10^−6^, q=0.04) was most significantly correlated with LINE-1 expression, pointing to activation of S phase checkpoint through ATM-MRN-SMC (8 other sites were more strongly correlated, but their functional significance is unknown). As part of the MRN complex, RAD50 is recruited to double strand breaks (DSB), where it is phosphorylated at S635 by ATM in response to ionizing radiation (IR). RAD50-S635 phosphorylation is necessary for ATM phosphorylation of the cohesin subunit SMC1A/SMC1 (Gatei et al., 2011), which in turn is necessary for the activation of S phase checkpoint in response to IR (Kim et al., 2002; Yazdi et al., 2002) (pathway shown left side of figure 4C). Indeed, ORF1p is strongly correlated with both known ATM-targeted phosphorylation sites on SMC1A (p<0.003) (figure 4D) and with four additional [ST]Q (ATM consensus motif) cohesin core residues: two on SMC1A and two on SMC3 (all p<0.05). ATM phosphorylation of SMC1A also requires BRCA1 and NBN/NBS1 (Kitagawa et al., 2004; Yazdi et al., 2002). At p<0.03, we find a correlation between ORF1p and ATM-targeted phosphosites on both proteins: BRCA1-S1545 (S1524 in uniprot) (M. Gatei et al., 2001) and NBN-S343 (Lim et al., 2000) (figure 4D). LINE-1 expression is not correlated with substrate abundance in any of these cases, consistent with the belief that these correlations are indicative of increased signaling (figure 4E, table S2). Despite this strong evidence for an association between LINE-1 expression and ATM signaling, we did not observe a significant correlation between ORF1p and ATM-S1981 autophosphorylation (ρ=0.14, p=0.29). ATM-S1981 is a popular measure of ATM activation (Bakkenist & Kastan, 2003), although it may not be essential (Pellegrini et al., 2006). We did however find enrichment for ATM activity using Kinase Set Enrichment Analysis (KSEA). The mean ORF1p/phosphositeplus ATM site correlation was 0.13 (p=7.8×10^−5^, figure 3F). Further supporting an interaction between LINE-1 and replication stress, we find that LINE-1 expression is correlated with all 7 core minichromosome maintenance complex (MCM) subunits at p<0.05. MCM is an essential helicase for DNA replication and has been shown to interact with LINE-1(Mita et al., 2018).

We next wanted to know whether there is also correlation between LINE-1 expression and the ATR-CHEK1 S phase checkpoint pathway. In response to replication stress, the RAD9A-HUS1-RAD1 (9-1-1) complex recruits ATR, which activates S phase checkpoint by phosphorylating CHEK1/CHK1 (right pathway of figure 4C) (Parrilla-Castellar et al., 2004). We find a strong correlation between ORF1p and ATR-T1989 autophosphorylation (ρ=0.37, p=0.0025). Given the overlapping motifs of ATM and ATR and the presence of a significant correlation with ATR but not ATM autophosphorylation, the MRN/SMC sites described above may be phosphorylated by ATR rather than ATM in this context. However, our analysis of LINE-1 expression in breast cancer (next section), and analyses by others (Coufal et al., 2011; Gasior et al., 2006), point to a role for ATM in the response to LINE-1. We were not able to assay ATR phosphorylation of CHEK1 due to a lack of phosphosite detection, but we did not find an enrichment in the phosphorylation of CHEK1 targets (figure 4F). Together this indicates that while there is an association between LINE-1 and ATR-CHEK1 activation, ATM-MRN-SMC is the primary checkpoint signalling that is correlated with LINE-1 expression in endometrial cancer.

Dou et al. and the CPTAC consortium indicated cell cycle dysregulation / checkpoint loss as a potential factor contributing to CNV in endometrial cancer. We found strong correlation between ORF1p and the cyclins E1, A2 and B1 (all ρ>0.4 and p<2.0×10^−4^) as well as enrichment for CDK1 and CDK2 phospho-targets (figure 5A-C), indicating that LINE-1 expression may be associated with cell cycle progression rather than successful checkpoint activation. Cyclin levels, however, do not appear to be driving global CNV independent of their correlation with LINE-1. Using partial correlation, we find that the ORF1p/CNV correlation remains significant after accounting for cyclin levels, but that these cyclins are not correlated with CNV after taking ORF1p levels into account (figure 5D).

**Figure 5:**
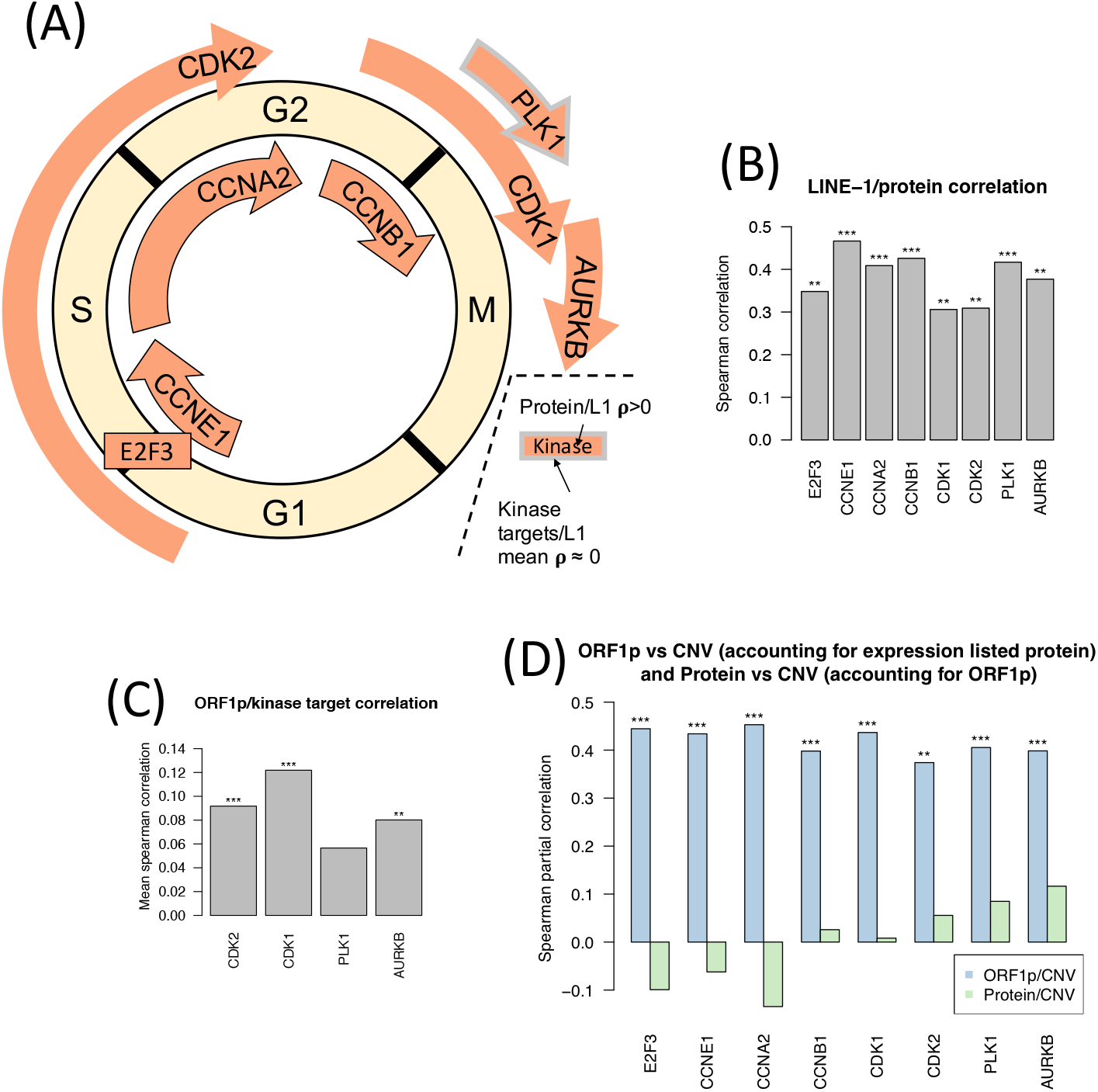
LINE-1/cell cycle progression correlations in endometrial cancer. (A) Drivers of cell cycle progression correlated with ORF1p, all at p<0.01. Kinases are outlined in red if there is significant enrichment for correlation between ORF1p and kinase targets, and gray if not. (B) Spearman correlation values for proteins shown in part A. *:0.01<p<0.05,**:0.001<p<0.01, ***:p<0.001. (C) KSEA for kinases shown in part A. (D) Partial correlation between ORF1p and CNV accounting for the level of these proteins (separately) and partial correlation between these proteins and CNV accounting for ORF1p levels. Correlation between ORF1p and CNV does not depend on these proteins, but correlation between these proteins and CNV does depend on ORF1p.

### In p53 wild type breast cancer, LINE-1 expression is associated with ATM deficiency, the ATR-CHEK1 signalling, NHEJ machinery and CNV

Our analysis of LINE-1 in endometrial cancer, is consistent with a model in which LINE-1 expression is associated with improperly repaired DSBs, likely occurring during S phase and possibly caused by replication fork collapse. We then wanted to know if this model is supported in other cancer types. In breast cancer, we found a weak correlation between LINE-1 expression and CNV (ρ=0.25, p=0.015). In the PAM50 breast cancer classification (Parker et al., 2009), the basal subtype, which has significant overlap with the triple negative classification, bears some similarities to the CNV high endometrial cancer subtype, including prevalent p53 mutation, elevated CNV and persistent DDR signaling (Krug et al., 2020). However, we did not find ORF1p to be elevated in basal breast cancer (figure 6A), despite finding that on average ORF1p levels are about 50% higher in p53 mutant breast cancers (p=0.011). Instead, we found that the LINE-1/CNV correlation in breast cancer is specific to p53 wild type tumors (ρ=0.32, p=0.017, figure 6B). We therefore set aside tumors with p53 mutations and focused only on the 57 breast tumors where ORF1p was measured, but no p53 mutation was detected.

**Figure 6:**
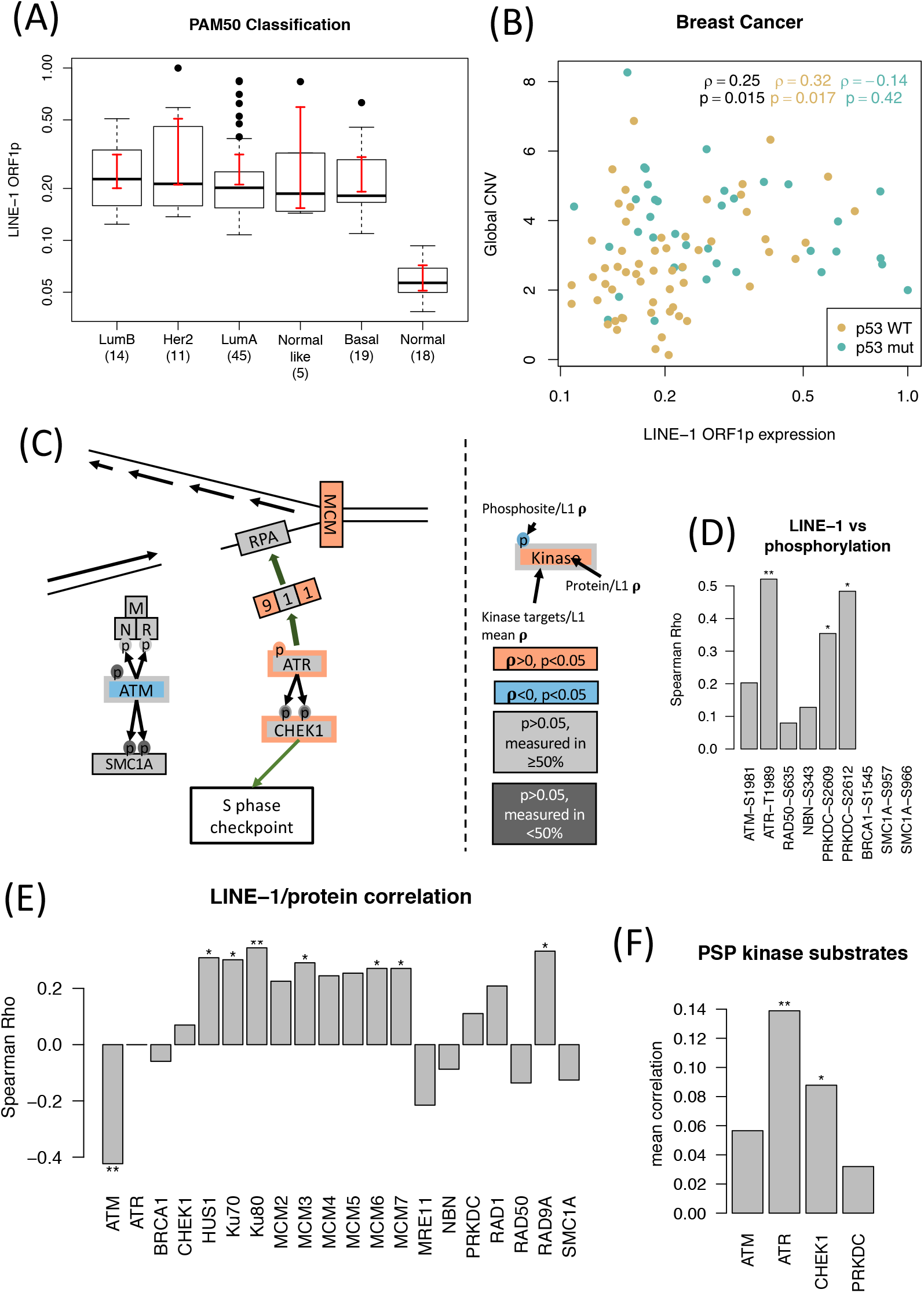
LINE/DDR correlation in p53 wild type breast cancer. (A) ORF1p quantification by PAM50 subtype. LINE-1 expression is universal across subtypes. (B) ORF1p vs global CNV. Despite ORF1p expression being on average higher in p53 mutant cancers, the LINE-1/CNV correlation is specific to p53 wild type. (C) A candidate model to explain the observed LINE-1/DDR correlations. At the top is a broken replication fork, on the left is the ATM-MRN-SMC pathway and on the right is the ATR-CHEK1 pathway. A protein or phosphosite filled in red indicates positive correlation at p<0.05, while blue indicates significant negative correlation, gray indicates no significant correlation, with dark gray indicating at least 50% missing values. Kinases are outlined in red if there is significant enrichment for correlation between ORF1p and kinase targets, and gray if not. Big green arrows indicate recruitment; small green arrows indicate downstream effects. (D) Spearman correlation between ORF1p and phosphosites pictured in C. *:0.01<p<0.05, **:0.001<p<0.01, ***:p<0.001. (E) As in D, but for ORF1p/protein correlation. (F) KSEA for correlation between ORF1p and kinase targets.

In the p53 wild type breast cancers, we did not find evidence for a correlation between LINE-1 and the ATM-MRN-SMC pathway (figure 6C, left pathway). Phosphosites on ATM, MRN, SMC1A and BRCA1 were either not detected or not correlated (figure 6D). This led us to ask whether there is some deficiency preventing activation of ATM-MRN-SMC in these cancers. Indeed, we found a strong anti-correlation between ORF1p and ATM protein level (ρ=−0.42, p=0.0011, figure 6E). Because ATM phosphorylates p53 and MDM2, leading to p53 stabilization and activation (Khosravi et al., 1999), this deficiency may explain how these cancers are able to elude the toxicity associated with high LINE-1 expression in cells with intact p53 (Ardeljan, Steranka, et al., 2020; Haoudi et al., 2004).

Cells with ATM deficiency would need to rely on other pathways to elicit checkpoint and repair DNA damage. We find a strong correlation between ORF1p and ATR-T1989 autophosphorylation (ρ=0.52, p=0.003) and enrichment for correlation with ATR (p=0.004) and with CHEK1 (p=0.01) phospho-targets (figure 6F). As in endometrial cancer, we could not assay ATR phosphorylation of CHEK1 as it was only detected in 5 tumors. Still, these correlations with measures of ATR/CHEK1 activation indicate that LINE-1 expressing, ATM deficient tumors are able to activate S phase checkpoint by relying on the ATR-CHEK1 pathway (figure 6C, right pathway).

An ATM deficiency may also indicate homologous recombination (HR) repair deficiency, forcing these tumors to rely on NHEJ (Morrison et al., 2000). Indeed, we see correlation between ORF1p and PRKDC autophosphorylation at T2609 and S2612 (figure 6D), and between ORF1p and NHEJ dimer subunits XRCC6/Ku70 and XRCC5/Ku80 (figure 6E) at the protein level (all p<0.04), but we do not see evidence for enrichment for correlation between LINE-1 expression and PRKDC targeted phosphosites (figure 6F). NHEJ, which is characterized as being more error prone than HR, is especially error prone when single ended DSB occur at collapsed replication forks (Balmus et al., 2019). Thus over reliance on NHEJ may promote CNV in LINE-1 expressing tumors. Correlations with the ATR-CHK1 pathway and NHEJ machinery remain significant after using partial correlation to account for ATM protein level (figure S3A), indicating that they are not merely a response to ATM deficiency.

Unlike in endometrial cancer, we did not see correlation between ORF1p and cyclins (except B2) or cell cycle related kinases such as CDK1/2, PLK1 and AURKB (figure S3B). However, we did find a weak (mean ρ<0.04), but significant (p<0.002) enrichment for correlation between LINE-1 expression and CDK1/2 phosphosites.

### In ovarian cancer LINE-1 expression is not associated with DDR phosphorylation but correlates with expression of PARP1 and proteins driving cell cycle progression

We next turned our attention to whether similar correlations between LINE-1 expression, CNV, DDR and cell cycle are present in ovarian cancer. While, we did observe a borderline correlation between LINE-1 ORF1p and CNV, it is driven by just four cases that have low LINE-1 and low CNV (figure 7A), and we did not identify significant correlation between LINE-1 expression and any of the above described DDR phosphosites. We did however find 80 proteins correlated with LINE-1 expression at FDR < 10% (table S2). These included four proteins known to interact with LINE-1: MOV10 (q=0.045), PABPC4 (q=0.032), PARP1 (q=0.056) and PURA (q=0.08) (Pizarro & Cristofari, 2016) (Figure 7C). MOV10 and PURA are both LINE-1 repressors (Taylor et al., 2018; Warkocki et al., 2018). PARP1 is a multifunctional DDR protein that is involved in the response to stalled replication forks (Liao et al., 2018). Its up-regulation could be a response to LINE-1 associated replication fork stalling.

**Figure 7:**
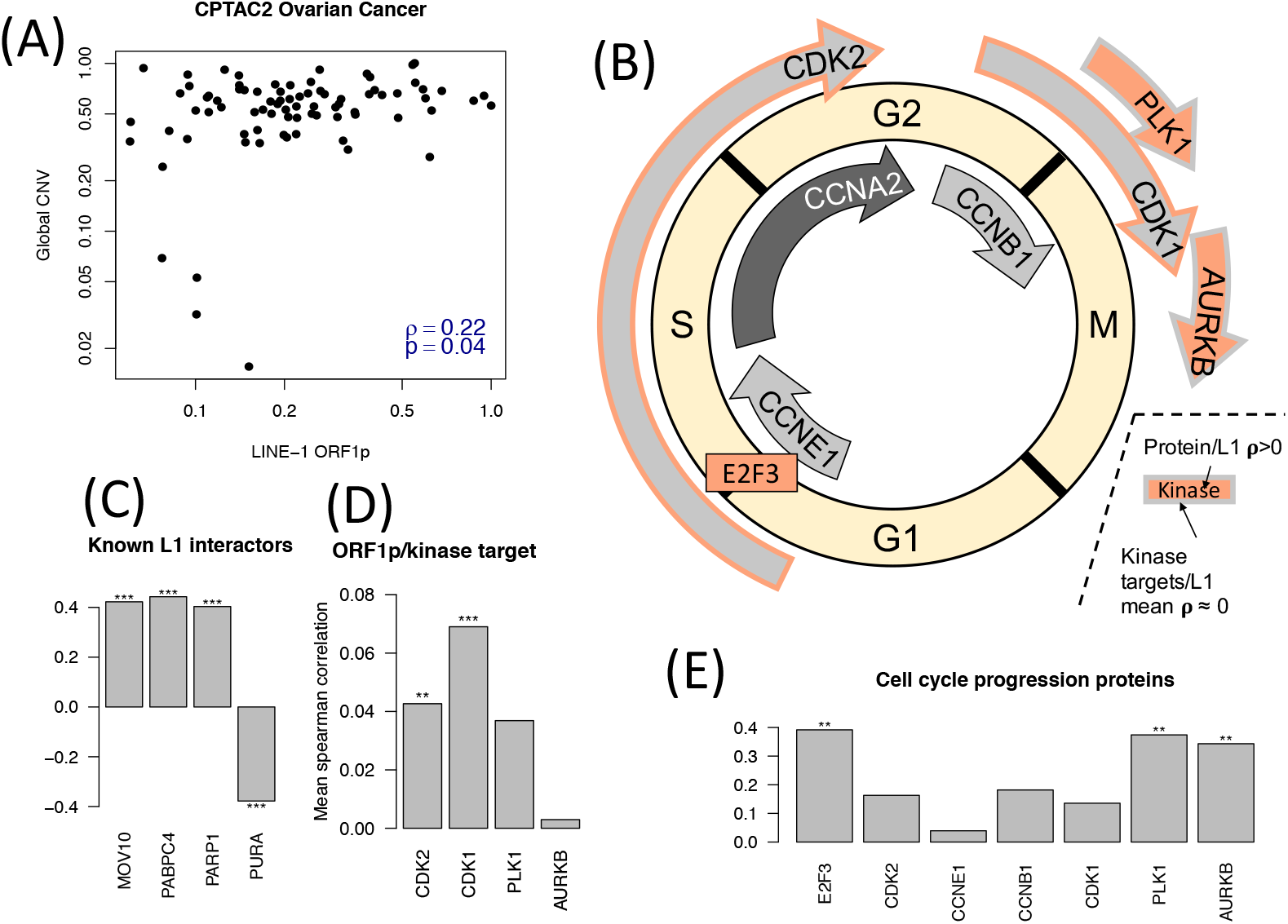
Analysis of LINE-1 in ovarian cancer. (A) ORF1p vs CNV. The correlation is significant at p<0.05, but depends on four cases with low LINE-1 and low CNV. (B) Correlation between LINE-1 and proteins that are known to interact with LINE-1 (all at q<0.1). *:0.01<p<0.05,**:0.001<p<0.01, ***:p<0.001. (C) Correlation between ORF1p and drivers of cell cycle progression. Red interior indicates positive correlation at p<0.05; insignificant correlations are in gray; CCNA2 correlation could not be clearly determined due to limited detection. Kinases are outlined in red if there is significant enrichment for correlation between ORF1p and kinase targets, and gray if not. (D) KSEA for kinases shown in part B. (E) ORF1p/Protein Spearman correlations for proteins shown in part B.

Next, we performed gene set enrichment analysis (GSEA), looking for Reactome (Jassal et al., 2020) gene sets enriched for correlation with ORF1p. “DNA replication” was the most enriched category after normalization (NES=2.74, p,q<10^−4^), with the G1/S transition driving transcription factor E2F3 being the most strongly correlated gene in the set (ρ=0.39, p=0.0068). Among the sets tested, “activation of ATR in response to replication stress” had the highest pre-normalization enrichment score (ES=0.75, p,q<10^−4^). This includes correlation with ORF1p at p<0.05 for 3 Replication Factor C complex (RFC) members, 2 Origin Recognition Complex (ORC) members, RAD9A and RPA1. Given the strong correlation between ORF1p and the cell cycle driving transcription factor E2F3, we wondered whether any of the other cell cycle driving ORF1p correlations identified in endometrial cancer are also present in ovarian cancer. We did not find a significant correlation with protein levels of cyclins E1, A2, and B1, but we did find significant ORF1p correlations for PLK1 (ρ=0.37, p=0.001) and AURKB (ρ=0.34, p=0.0025) protein levels and CDK1 and 2 phosphotargets (Figure 7B, D-E).

## DISCUSSION

We have used phosphoproteomic data to identify correlations between LINE-1 expression and S phase checkpoint signaling. This indicates that recent studies showing that LINE-1 overexpression leads to DNA replication conflicts are relevant to human tumors. We also identified strong LINE-1 RNA/protein correlations that exceed the RNA/protein correlation for most host genes as well as correlations between LINE-1 expression and drivers of cell cycle progression that may indicate changes to cell cycle in response to chronic DNA damage or may indicate faster growth in LINE-1 expressing tumors.

One might expect that because of the challenges associated with quantifying the expression of highly repetitive elements such as LINE-1, the LINE-1 RNA/protein correlation would be weaker than those for most host genes. The very high degree of correlation that we observe would then be surprising. However, ORF1p binds in cis to its own RNA (Wei et al., 2001), likely protecting it from degradation. If naked LINE-1 RNAs are highly susceptible to degradation, most LINE-1 RNA would be present in complex with ORF1p, explaining the elevated RNA/protein correlation. Correlation between LINE-1 RNA and ORF2p or between ORF1p and ORF2p is less clear. Endogenous ORF2p is extremely difficult to measure (Ardeljan, Wang, et al., 2020) and was at best sparsely observed in our analysis, but we did find that correlation LINE-1 ORF1p and somatic insertions are correlated in endometrial cancer. Because ORF2p is critical to retrotransposition, this suggests that it is also correlated with somatic insertions and thus with ORF1p, at least in endometrial cancer. Translational regulation of ORF2p is well reported (Alisch et al., 2006; Mita et al., 2018; Taylor et al., 2013) and WGS was limited, so such a correlation may not be universal. It is also possible to have ORF1p expression from loci that lack an intact ORF2. However, we find ORF1p to be similarly correlated with both intact and ORF1 only RNA (figure S1).

Our finding that LINE-1 expression is correlated with S phase checkpoint signaling in endometrial and p53 wildtype breast cancer provides the first in vivo support for the hypothesis that LINE-1 expression causes replication stress and persistent DNA damage. In endometrial cancer, we see correlation between LINE-1 and ATM-MRN-SMC signaling. This pathway is activated when cells are exposed to ionizing radiation during S phase (Magtouf Gatei et al., 2011; Luo et al., 2008; Yazdi et al., 2002), suggesting that LINE-1 expression is correlated with ds breaks occurring during S phase. Such breaks could result from the collapse of replication forks after collision with LINE-1 retrotransposition intermediates. In p53 wild-type breast cancer, we saw a correlation between LINE-1 expression and ATR-CHEK1 signaling. This would at first seem to suggest a different type of DNA damage as ATR is associated with stalled replication forks rather than ds breaks (Saldivar et al., 2017). However, LINE-1 expression is associated with ATM deficiency in these tumors and ATR can be activated at resected ds breaks (Smith et al., 2010), so ATR-CHEK1 may be activated as an alternative to ATM-MRN-SMC1A. We did not identify clear correlations between LINE-1 expression and DDR signaling in p53 mutant breast cancers, despite the presence of persistent DDR signaling in many of these tumors. It is likely that, in these tumors, any effect of LINE-1 on DNA damage is overwhelmed by other factors. We also did not identify LINE-1 expression / DDR phosphorylation in ovarian cancer, but we did find a correlation with PARP1 expression, hinting at a DNA damage connection. In fact, we did not find any LINE-1 expression / phosphosite correlations that were significant after multiple hypothesis correction. Ovarian cancer had the highest level of LINE-1 RNA in our study, so it may be that LINE-1 expression is saturated in ovarian cancer, limiting the efficacy of our methods. Finally, in colon cancer, we found our LINE-1 ORF1p quantification to be highly correlated with tumor purity (as measured by ESTIMATE (Yoshihara et al., 2013)). We could not discern whether this is a biologically relevant result or whether the LINE-1 signal is simply diluted in low purity tumors, so we provided only limited analysis of colon cancer in this study.

Finally, we find that LINE-1 expression is correlated with several drivers of cell cycle progression. In both endometrial and ovarian cancer, we find correlation between ORF1p and the G1 to S promoting transcription factor E2F3 and between ORF1p and the G2 to M promoting kinase PLK1. We also find evidence for increased cyclin dependent kinase (CDK) activity based on kinase set enrichment analysis. Because this data is from bulk tissue, we cannot be certain whether these correlations reflect an up-regulation of cell-cycle progression signaling or stalling and accumulation at particular cell cycle stages. Cell cycle stalling and accumulation is consistent with the hypothesis that LINE-1 expression leads to persistent replication stress and DNA damage. Challenges to DNA replication could explain increases in signaling and expression associated with cells in S or G2 phase. At the same time, repeated studies have found that LINE-1 de-repression is associated with poor prognosis (not analyzed in this study because of lack of follow up data) in many cancer types (reviewed in (Ardeljan et al., 2017)). It may therefore be that these correlations do reflect a more rapid transit through cell cycle, perhaps as an adaptation to the stress caused by LINE-1 expression or perhaps reflecting some unknown mechanism. Consistent with this, correlations with M phase activity such as cyclin B-CDK1 are not easily explained by replication stress and S phase checkpoint activation. However, DNA synthesis can continue into mitosis (Minocherhomji et al., 2015), so further experiments are needed to understand the functional meaning of these correlations.

Our study shows that LINE-1 expression can be consistently measured in large multi-omic cancer datasets. We then leverage this to find in vivo evidence that LINE-1 expression is correlated with DNA damage occurring in the S phase of cell cycle and to identify correlations between LINE-1 expression and markers of cell cycle progression. In endometrial cancer, there are 73 phosphosites correlated with LINE-1 expression at q<0.1 and more than 5000 at p<0.05, most of which are unstudied. As more is known about the role of individual phosphosites, it may be possible to return to these analyses and find additional pathways that are correlated with LINE-1 expression. In particular, LINE-1 expression is associated with an interferon response (Ardeljan, Steranka, et al., 2020; Cecco et al., 2019; Yu et al., 2015) that we were not able to elucidate here.

## SUPPLEMENTARY FIGURE CAPTIONS

**Figure S1.**
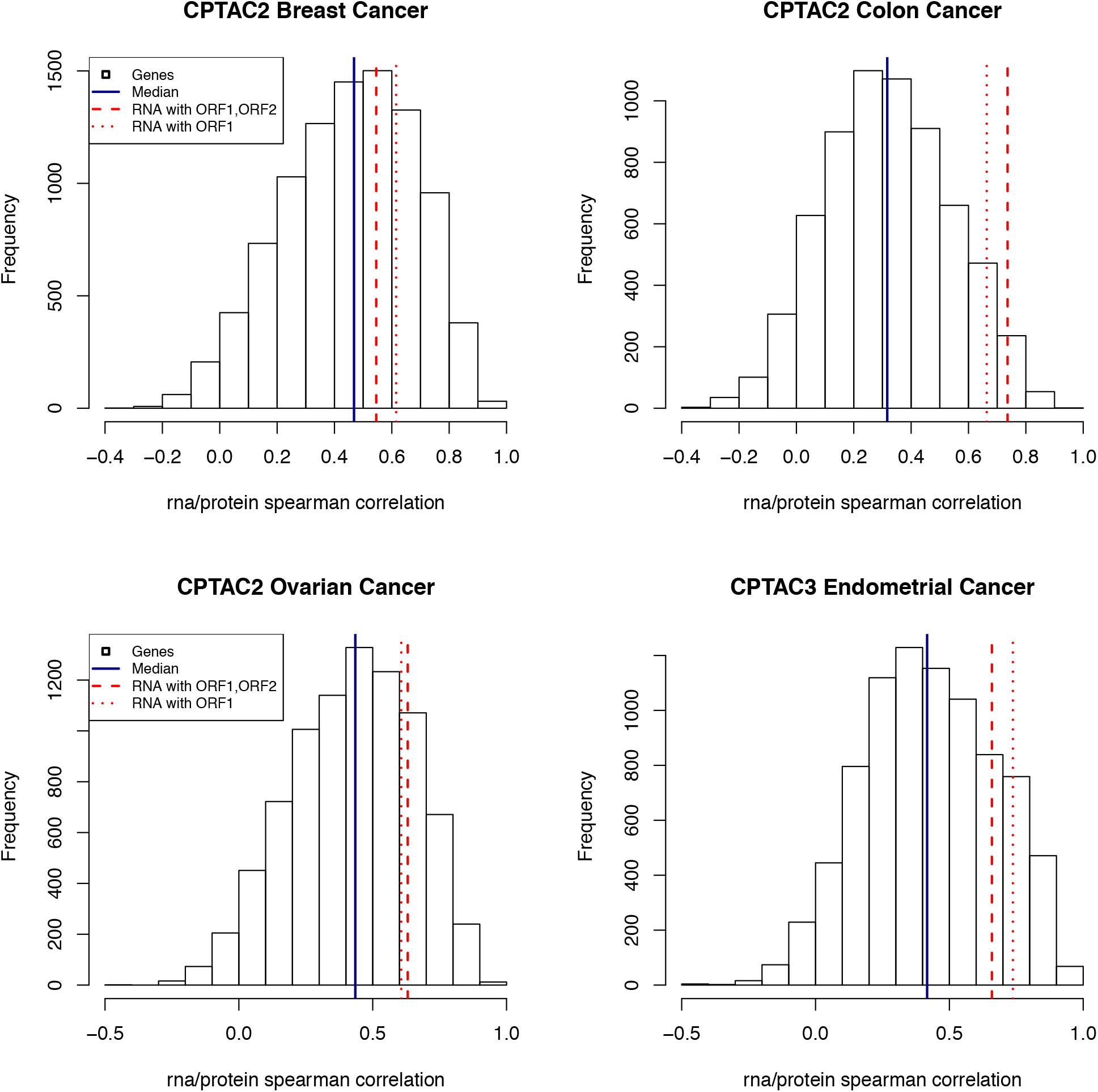
Histogram of RNA/protein correlations for all human proteins measured in at least half of samples, overlayed with the correlation between LINE-1 ORF1p and LINE-1 RNA assigned to intact loci (dashed red lines) or to loci with at least ORF1 intact (dotted red lines). (A) Breast cancer, (B) ovarian cancer, (C) colon cancer, (D) endometrial cancer.

**Figure S2.**
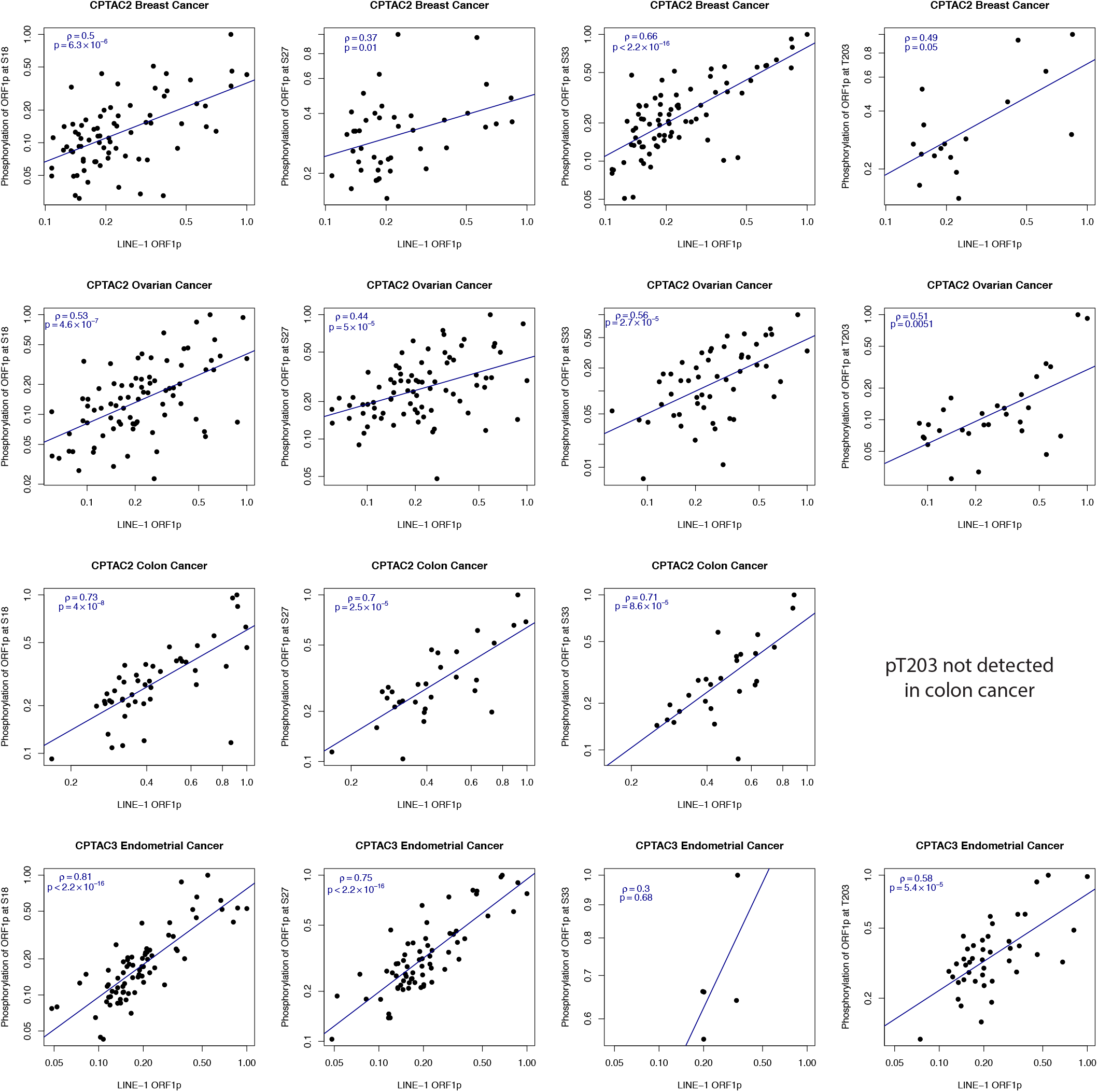
Correlation between ORF1p expression and phosphorylation at the 4 identified phosphosites. Rows correspond to cancer types: breast, ovarian, colon and endometrial. Columns correspond to phosphosites: S18, S27, S33, T203.

**Figure S3.**
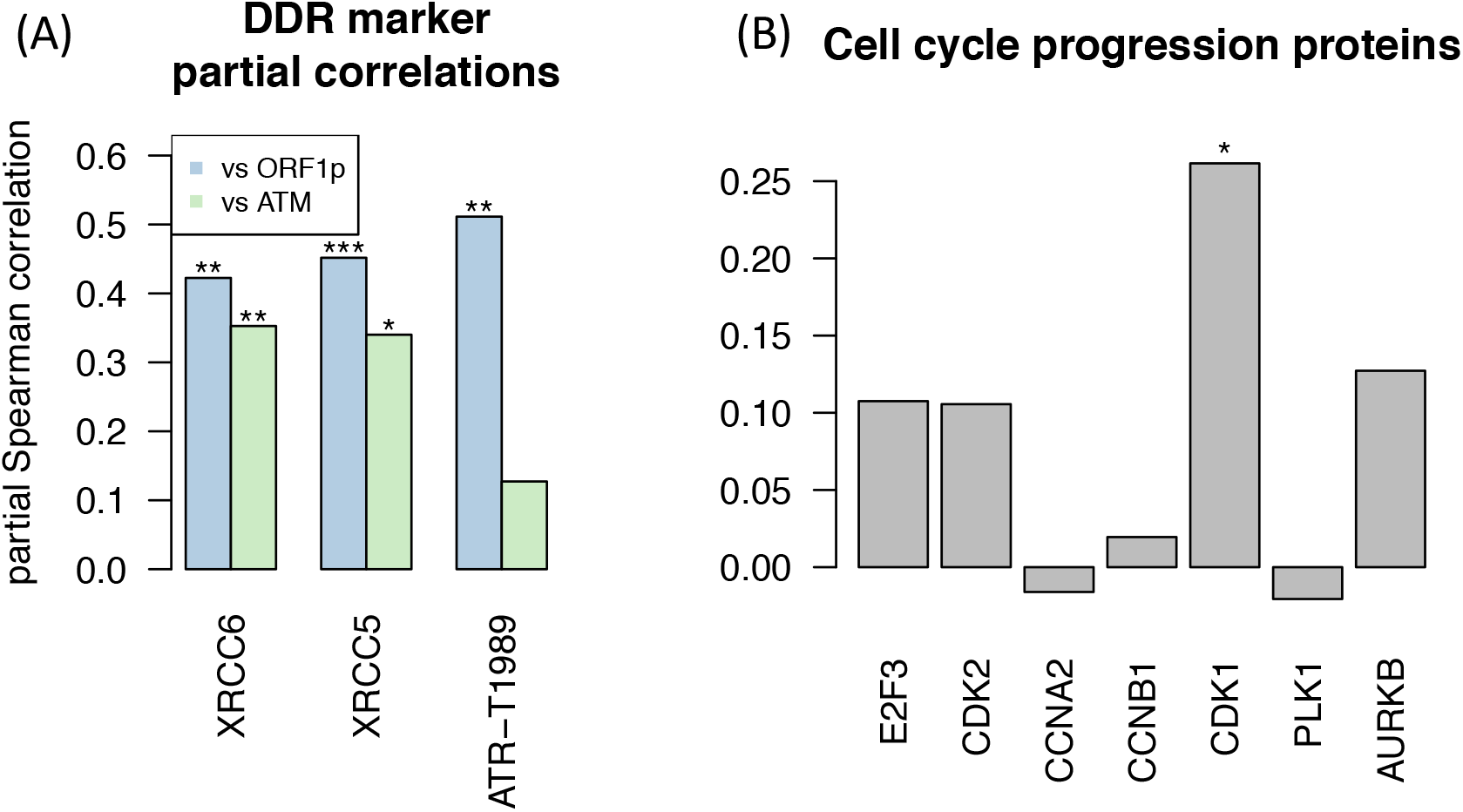
Additional breast cancer plots. (A) Partial Spearman correlation between ORF1p and DDR protein/phosphosite, accounting for ATM, and partial Spearman correlation between ATM and DDR proteins/phosphosites, accounting for ORF1p. ATM alone deficiency does not explain ORF1p/Ku and ORF1p/ATR-T1989 correlations.

**Figure S4.**
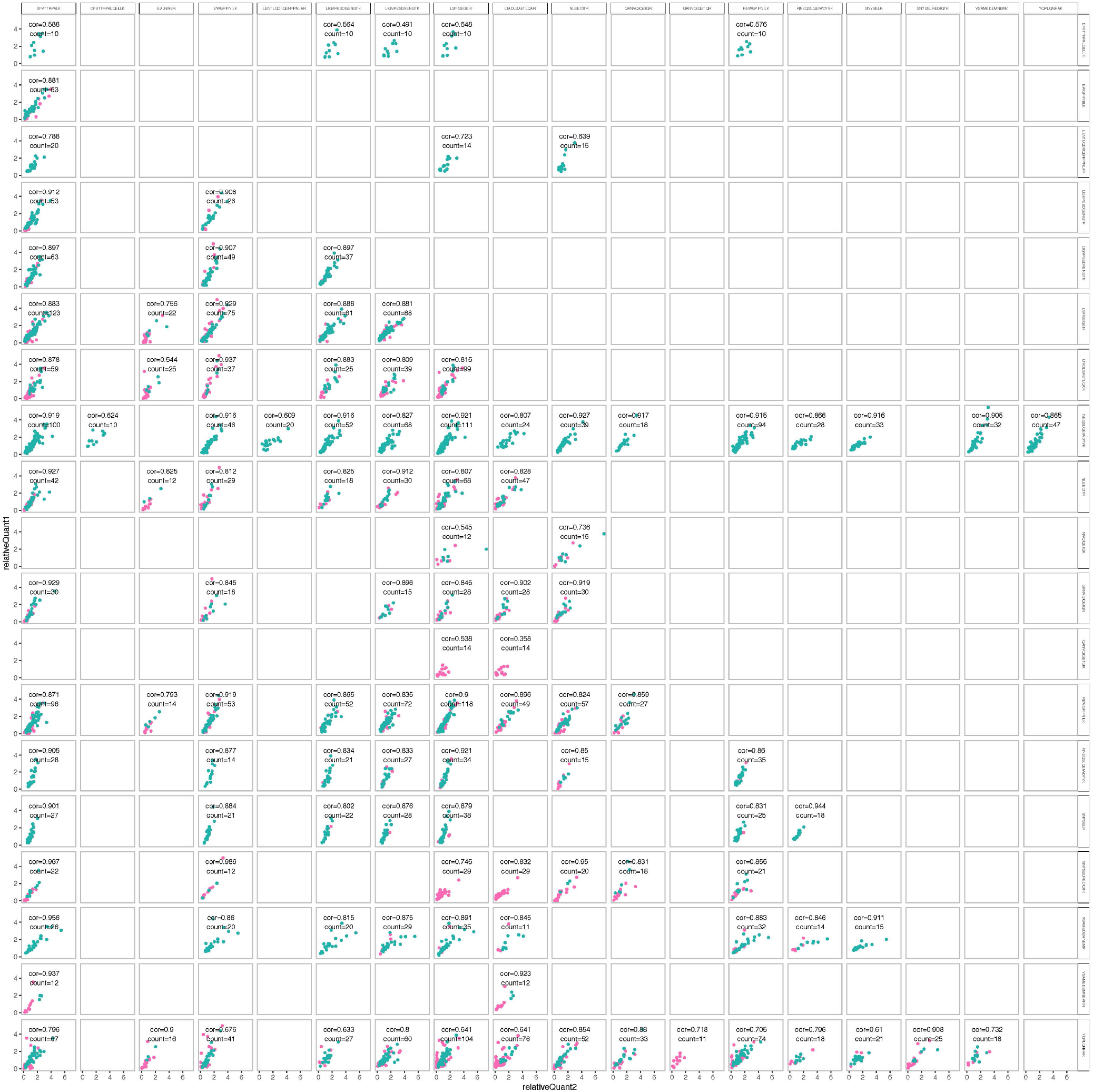
Spearman correlations between the relative quantification of ORF1p peptides in CPTAC2 retrospective data. Breast cancer samples are in pink; ovarian samples are teal. Peptides used for quantification correlate with two other ORF1p peptides at ρ>0.6.

## Supplementary table captions

**Table S1**. Quantifications of LINE-1 RNA, ORF1p, insertion and phosphorylation. Tumor samples only.

**Table S2**. Correlation between ORF1p and host protein levels. Tumor samples only.

**Table S3**. Correlation between ORF1p and phosphosites. Tumor samples only.

## STAR METHODS

### Quantification of LINE-1 RNA

LINE-1 RNA was quantified from available RNA-seq data using an implementation of L1EM (McKerrow & Fenyö, 2020) on the Cancer Genome Cloud. L1EM employs the expectation maximization algorithm to estimate locus specific gene expression and to separate proper LINE-1 expression from passive co-transcription that includes LINE-1 RNA, but does not support retrotransposition. For the intact LINE-1 RNA quantifications, full length LINE-1 loci with no stop codon in either ORF1 or ORF2 were considered expressed if at least 2 read pairs per million (FPM) were assigned to that locus and less than 10% of the RNA assigned to that locus was estimated to be passive co-transcription. Total intact LINE-1 RNA expression was estimated by adding together the FPM values for each such locus. Full length loci with stop codons in ORF2 but not ORF1 were also included to generate an ORF1 RNA expression estimate. Samples were excluded if no locus was detected at 2 FPM as this may be due to data quality rather than a lack of LINE-1 RNA. Specifically, ovarian cancer samples that failed LINE-1 RNA quantification did not have lower ORF1p quantifications.

### Quantification of LINE-1 ORF1p

LINE-1 ORF1p was quantified from isobaric labelled MS/MS proteomics data using a set of 20 proteotypic ORF1p peptides generated from analysis of older CPTAC breast (Mertins et al., 2016) and ovarian cancer (Zhang et al., 2016) data. X!tandem (Craig & Beavis, 2004) was used to search mass spectra against a combined database that includes both the standard ensembl human proteome and in silico translations of intact LINE-1 open reading frames in the human reference genome. Oxidation of methionine (+15.994915@M) was included as a potential modification. Carbamidomethylation of cysteine (+57.022@C) was set as a fixed modification in addition to modifications appropriate to the particular isobaric labelling used (+144.102063@[,+144.102063@K for iTRAQ4 and +229.162932@[, +229.162932@K for TMT10). Peptides were quantified by calculating the log ratio between the reporter intensity for each sample and the reporter intensity of an internal control. Peptide/peptide spearman correlations were calculated and peptides that had a correlation of at least 0.6 with two other peptides were retained (see figure S4). This led to the following list: DFVTTRPALK, EALNMER, EWGPIFNILK, LIGVPESDGENGTK, LIGVPESDVENGTK, LSFISEGEIK, LTADLSAETLQAR, NLEECITR, NVQIQEIQR, QANVQIQEIQR, REWGPIFNILK, RNEQSLQEIWDYVK, SNYSELR, SNYSELREDIQTK, VSAMEDEMNEMK, YQPLQNHAK, DFVTTRPALQELLK, LENTLQDIIQENFPNLAR, VSAMEDEMNEMKR, and NEQSLQEIWDYVK.

For the actual peptide quantifications, the x!tandem search and peptide quantifications were performed as above. If a peptide were detected multiple times in the same sample, the median across peptide spectral matches was used. Then to get a protein quantification, the median was taken across all peptides in the preceding list that were identified in that sample. Finally, the quantification was translated by the median log ratio for all human proteins to account for variation in the size of the input sample. This pipeline was implemented and executed as a workflow on the Cancer Genome Cloud.

### Quantification of LINE-1 ORF1p phosphorylation

ORF1p phosphopeptides were identified using the above x!tandem search strategy with the addition of phosphorylation at serine, threonine and tyrosine (79.966331@S, 79.966331@T, 79.966331@Y) as potential modifications. Identified ORF1p phosphopeptides were aligned to the uniprot LINE-1 ORF1p sequence (L1RE1) using the pairwise2 method in Biopython to identify the phosphorylation coordinate. Phosphopeptides were quantified as above using the log ratio between the sample reporter intensity and the reference reporter intensity. If multiple phosphopeptide spectral matches indicated the same phosphosite, the median was taken.

### Quantification of high confidence LINE-1 somatic insertions

Somatic insertions were identified using the Mobile Element Locator Tool (MELT) (Gardner et al., 2017) on the Cancer Genome Cloud. MELT was run on each pair of matched cancer/normal WGS datasets. Only insertions with the greatest evidence (ASSESS=5) were considered. To be considered a high confidence somatic insertion, the insertion had to be called heterozygous in cancer and homozygous absent in normal, with the log likelihood of both of these genotypes being at least 10 greater than the log likelihood of the next most probable genotype.

### Estimation of global CNV

For breast and endometrial cancer, global CNV scores were provided by the respective CPTAC working groups (Krug et al., 2020; (Dou et al., 2020). For colon cancer, the CNV score was recalculated according to the description provided (Vasaikar et al., 2019). For ovarian cancer, a global CNV score was calculated from gene level CNV provided by the CPTAC working group (McDermott et al., 2020). First, any gene overlapping another gene with a smaller leftmost coordinate was removed. Then, each remaining gene coordinate range was extended in the plus direction to reach the start of the next gene. The global CNV score was then a linear combination of the absolute value of the gene level CNV with the length of the extended gene ranges.

### Enrichment Analysis

For gene set enrichment analysis (GSEA), Spearman correlations were calculated between our LINE-1 ORF1p quantification and the log normalized quantification for each identified protein identified in at least half of samples. Analysis was then performed using the GSEAPreranked option in GSEA 4.0.0. Protein searches were performed against Reactome gene sets (Jassal et al., 2020).

For kinase set enrichment analysis (KSEA), we calculated partial Spearman correlation between ORF1p and each phosphosite that was identified in at least half samples, accounting for the substrate protein quantification. Enrichment was calculated using the KSEA web app (Wiredja et al., 2017).

### Statistical analysis

Statistical analysis was performed in R. Spearman correlation was calculated using the cor.test function in the standard stats package. Partial correlation was calculated using pcortest in the ppcor package. The Benjamini-Hochberg (BH) method was used to calculate FDR/q value when considering multiple hypothesis tests. Uncorrected p values were used when literature or analysis of another cancer type pointed to potential involvement of a specific gene/protein/phosphosite.

## Data and Code Availability

Raw and processed proteomic and phosphoproteomic data can be found at the CPTAC portal (https://cptac-data-portal.georgetown.edu/cptacPublic/) or the Proteomic Data Commons (https://pdc.cancer.gov/pdc/). Transcriptomic and genomic data for CPTAC 3 (endometrial and kidney cancers) can be found in the Genomic Data Commons (https://gdc.cancer.gov/). Additionally data for each cancer type can be found in the corresponding publication: colon (Vasaikar et al., 2019), breast (Krug et al., 2020), ovarian (McDermott et al., 2020), endometrial (Dou et al., 2020), and kidney (Clark et al., 2019). L1EM code can be found at https://github.com/FenyoLab/L1EM. CGC implementations of L1EM and ORF1p quantification are available on request (please provide a CGC username).

## Funding Acknowledgment

This project has been funded in whole or in part with Federal funds from the National Cancer Institute, National Institutes of Health, under Contract No. HHSN261200800001E. The content of this publication does not necessarily reflect the views or policies of the Department of Health and Human Services, nor does mention of trade names, commercial products, or organizations imply endorsement by the U.S. Government. Additional funding was provided by NIH grants P01AG051449 (NIA subcontracts to J.B. and D.F.), U24CA210972 (NCI to D.F.), and 1R21CA235521 (NCI to J.B.).

## Author Contributions

D.F. and X.W. conceived the project. X.W., with M.G. and S.C., developed and implemented LINE-1 protein quantification methods. W.M. performed cloud based analyses and data integration. W.M. led manuscript preparation with guidance on framing and significance from P.M., L.D., J.L., J.B., and D.F. provided project management and oversight.

## References

Krug, K., Jaehnig, E. J., Blumenberg, L., Karpova, A., Anurag, M., Miles, G., Mertins, P., Geffen Y., Tang, L. C., Heiman, D. I., Cao, S., Maruvka, Y., Lei, J. T., Huang, C., Kothadia, R. B., Colaprico, A., Wiznerowicz, M., Wyczalkowski, M. A., Thiagarajan, M., Kinsinger, C. R., Hiltke, T., Boja, E., Mesri, M., Robles, A. I., Rodriguez, H., Ding, L., Getz, G., Karl R. Clauser, K. R., Fenyö, D., Ruggles, R., Zhang, B., Mani, D. R., Satpathy, S., Carr, S. A., Ellis, M. J., Gillette, M. A., Clinical Proteomic Tumor Analysis Consortium. (2020). Proteogenomics redefines breast cancer biological subtype classifications and nominates therapeutic targets. Submitted to Cell.

Alisch, R. S., Garcia-Perez, J. L., Muotri, A. R., Gage, F. H., & Moran, J. V. (2006). Unconventional translation of mammalian LINE-1 retrotransposons. Genes & Development, 20(2), 210–224. https://doi.org/10.1101/gad.1380406

Ardeljan, D., Steranka, J. P., Liu, C., Li, Z., Taylor, M. S., Payer, L. M., Gorbounov, M., Sarnecki, J. S., Deshpande, V., Hruban, R. H., Boeke, J. D., Fenyö, D., Wu, P.-H., Smogorzewska, A., Holland, A. J., & Burns, K. H. (2020). Cell fitness screens reveal a conflict between LINE-1 retrotransposition and DNA replication. Nature Structural & Molecular Biology, 27(2), 168–178. https://doi.org/10.1038/s41594-020-0372-1

Ardeljan, D., Taylor, M. S., Ting, D. T., & Burns, K. H. (2017). The Human Long Interspersed Element-1 Retrotransposon: An Emerging Biomarker of Neoplasia. Clinical Chemistry, 63(4), 816–822. https://doi.org/10.1373/clinchem.2016.257444

Ardeljan, D., Wang, X., Oghbaie, M., Taylor, M. S., Husband, D., Deshpande, V., Steranka, J. P., Gorbounov, M., Yang, W. R., Sie, B., Larman, H. B., Jiang, H., Molloy, K. R., Altukhov, I., Li, Z., McKerrow, W., Fenyö, D., Burns, K. H., & LaCava, J. (2020). LINE-1 ORF2p expression is nearly imperceptible in human cancers. Mobile DNA, 11, 1. https://doi.org/10.1186/s13100-019-0191-2

Bakkenist, C. J., & Kastan, M. B. (2003). DNA damage activates ATM through intermolecular autophosphorylation and dimer dissociation. Nature, 421(6922), 499–506. https://doi.org/10.1038/nature01368

Balmus, G., Pilger, D., Coates, J., Demir, M., Sczaniecka-Clift, M., Barros, A. C., Woods, M., Fu, B., Yang, F., Chen, E., Ostermaier, M., Stankovic, T., Ponstingl, H., Herzog, M., Yusa, K., Martinez, F. M., Durant, S. T., Galanty, Y., Beli, P., … Jackson, S. P. (2019). ATM orchestrates the DNA-damage response to counter toxic non-homologous end-joining at broken replication forks. Nature Communications, 10(1), 87. https://doi.org/10.1038/s41467-018-07729-2

Belancio, V. P., Roy-Engel, A. M., Pochampally, R. R., & Deininger, P. (2010). Somatic expression of LINE-1 elements in human tissues. Nucleic Acids Research, 38(12), 3909–3922. https://doi.org/10.1093/nar/gkq132

Beraldi, R., Pittoggi, C., Sciamanna, I., Mattei, E., & Spadafora, C. (2006). Expression of LINE-1 retroposons is essential for murine preimplantation development. Molecular Reproduction and Development, 73(3), 279–287. https://doi.org/10.1002/mrd.20423

Cajuso, T., Sulo, P., Tanskanen, T., Katainen, R., Taira, A., Hänninen, U. A., Kondelin, J., Forsström, L., Välimäki, N., Aavikko, M., Kaasinen, E., Ristimäki, A., Koskensalo, S., Lepistö, A., Renkonen-Sinisalo, L., Seppälä, T., Kuopio, T., Böhm, J., Mecklin, J.-P., … Aaltonen, L. A. (2019). Retrotransposon insertions can initiate colorectal cancer and are associated with poor survival. Nature Communications, 10(1), 4022. https://doi.org/10.1038/s41467-019-11770-0

Cancer Genome Atlas Research Network, Kandoth, C., Schultz, N., Cherniack, A. D., Akbani, R., Liu, Y., Shen, H., Robertson, A. G., Pashtan, I., Shen, R., Benz, C. C., Yau, C., Laird, P. W., Ding, L., Zhang, W., Mills, G. B., Kucherlapati, R., Mardis, E. R., & Levine, D. A. (2013). Integrated genomic characterization of endometrial carcinoma. Nature, 497(7447), 67–73. https://doi.org/10.1038/nature12113

Cecco, M. D., Ito, T., Petrashen, A. P., Elias, A. E., Skvir, N. J., Criscione, S. W., Caligiana, A., Brocculi, G., Adney, E. M., Boeke, J. D., Le, O., Beauséjour, C., Ambati, J., Ambati, K., Simon, M., Seluanov, A., Gorbunova, V., Slagboom, P. E., Helfand, S. L., … Sedivy, J. M. (2019). L1 drives IFN in senescent cells and promotes age-associated inflammation. Nature, 566(7742), 73. https://doi.org/10.1038/s41586-018-0784-9

Clark, D. J., Dhanasekaran, S. M., Petralia, F., Pan, J., Song, X., Hu, Y., da Veiga Leprevost, F., Reva, B., Lih, T.-S. M., Chang, H.-Y., Ma, W., Huang, C., Ricketts, C. J., Chen, L., Krek, A., Li, Y., Rykunov, D., Li, Q. K., Chen, L. S., … Clinical Proteomic Tumor Analysis Consortium. (2019). Integrated Proteogenomic Characterization of Clear Cell Renal Cell Carcinoma. Cell, 179(4), 964–983.e31. https://doi.org/10.1016/j.cell.2019.10.007

Cook, P. R., Jones, C. E., & Furano, A. V. (2015). Phosphorylation of ORF1p is required for L1 retrotransposition. Proceedings of the National Academy of Sciences of the United States of America, 112(14), 4298–4303. https://doi.org/10.1073/pnas.1416869112

Coufal, N. G., Garcia-Perez, J. L., Peng, G. E., Marchetto, M. C. N., Muotri, A. R., Mu, Y., Carson, C. T., Macia, A., Moran, J. V., & Gage, F. H. (2011). Ataxia telangiectasia mutated (ATM) modulates long interspersed element-1 (L1) retrotransposition in human neural stem cells. Proceedings of the National Academy of Sciences of the United States of America, 108(51), 20382–20387. https://doi.org/10.1073/pnas.1100273108

Craig, R., & Beavis, R. C. (2004). TANDEM: Matching proteins with tandem mass spectra. Bioinformatics (Oxford, England), 20(9), 1466–1467. https://doi.org/10.1093/bioinformatics/bth092

Dou, Y., Kawaler, E. A., Cui Zhou, D., Gritsenko, M. A., Huang, C., Blumenberg, L., Karpova, A., Petyuk, V. A., Savage, S. R., Satpathy, S., Liu, W., Wu, Y., Tsai, C.-F., Wen, B., Li, Z., Cao, S., Moon, J., Shi, Z., Cornwell, M., … Clinical Proteomic Tumor Analysis Consortium. (2020). Proteogenomic Characterization of Endometrial Carcinoma. Cell, 180(4), 729–748.e26. https://doi.org/10.1016/j.cell.2020.01.026

Doucet, A. J., Wilusz, J. E., Miyoshi, T., Liu, Y., & Moran, J. V. (2015). A 3’ Poly(A) Tract Is Required for LINE-1 Retrotransposition. Molecular Cell, 60(5), 728–741. https://doi.org/10.1016/j.molcel.2015.10.012

Feng, Q., Moran, J. V., Kazazian, H. H., & Boeke, J. D. (1996). Human L1 retrotransposon encodes a conserved endonuclease required for retrotransposition. Cell, 87(5), 905–916.

Gaillard, H., García-Muse, T., & Aguilera, A. (2015). Replication stress and cancer. Nature Reviews. Cancer, 15(5), 276–289. https://doi.org/10.1038/nrc3916

Gardner, E. J., Lam, V. K., Harris, D. N., Chuang, N. T., Scott, E. C., Pittard, W. S., Mills, R. E., Consortium, 1000 Genomes Project, & Devine, S. E. (2017). The Mobile Element Locator Tool (MELT): Population-scale mobile element discovery and biology. Genome Research, gr.218032.116. https://doi.org/10.1101/gr.218032.116

Gasior, S. L., Wakeman, T. P., Xu, B., & Deininger, P. L. (2006). The Human LINE-1 Retrotransposon Creates DNA Double-strand Breaks. Journal of Molecular Biology, 357(5), 1383–1393. https://doi.org/10.1016/j.jmb.2006.01.089

Gatei, M., Zhou, B. B., Hobson, K., Scott, S., Young, D., & Khanna, K. K. (2001). Ataxia telangiectasia mutated (ATM) kinase and ATM and Rad3 related kinase mediate phosphorylation of Brca1 at distinct and overlapping sites. In vivo assessment using phospho-specific antibodies. The Journal of Biological Chemistry, 276(20), 17276–17280. https://doi.org/10.1074/jbc.M011681200

Gatei, Magtouf, Jakob, B., Chen, P., Kijas, A. W., Becherel, O. J., Gueven, N., Birrell, G., Lee, J.-H., Paull, T. T., Lerenthal, Y., Fazry, S., Taucher-Scholz, G., Kalb, R., Schindler, D., Waltes, R., Dörk, T., & Lavin, M. F. (2011). ATM protein-dependent phosphorylation of Rad50 protein regulates DNA repair and cell cycle control. The Journal of Biological Chemistry, 286(36), 31542–31556. https://doi.org/10.1074/jbc.M111.258152

Haoudi, A., Semmes, O. J., Mason, J. M., & Cannon, R. E. (2004). Retrotransposition-Competent Human LINE-1 Induces Apoptosis in Cancer Cells With Intact p53. Journal of Biomedicine & Biotechnology, 2004(4), 185–194. https://doi.org/10.1155/S1110724304403131

Helman, E., Lawrence, M. S., Stewart, C., Sougnez, C., Getz, G., & Meyerson, M. (2014). Somatic retrotransposition in human cancer revealed by whole-genome and exome sequencing. Genome Research, 24(7), 1053–1063. https://doi.org/10.1101/gr.163659.113

Hornbeck, P. V., Zhang, B., Murray, B., Kornhauser, J. M., Latham, V., & Skrzypek, E. (2015). PhosphoSitePlus, 2014: Mutations, PTMs and recalibrations. Nucleic Acids Research, 43(Database issue), D512–520. https://doi.org/10.1093/nar/gku1267

Jassal, B., Matthews, L., Viteri, G., Gong, C., Lorente, P., Fabregat, A., Sidiropoulos, K., Cook, J., Gillespie, M., Haw, R., Loney, F., May, B., Milacic, M., Rothfels, K., Sevilla, C., Shamovsky, V., Shorser, S., Varusai, T., Weiser, J., … D’Eustachio, P. (2020). The reactome pathway knowledgebase. Nucleic Acids Research, 48(D1), D498–D503. https://doi.org/10.1093/nar/gkz1031

Jung, H., Choi, J. K., & Lee, E. A. (2018). Immune signatures correlate with L1 retrotransposition in gastrointestinal cancers. Genome Research, 28(8), 1136–1146. https://doi.org/10.1101/gr.231837.117

Kano, H., Godoy, I., Courtney, C., Vetter, M. R., Gerton, G. L., Ostertag, E. M., & Kazazian, H. H. (2009). L1 retrotransposition occurs mainly in embryogenesis and creates somatic mosaicism. Genes & Development, 23(11), 1303–1312. https://doi.org/10.1101/gad.1803909

Khosravi, R., Maya, R., Gottlieb, T., Oren, M., Shiloh, Y., & Shkedy, D. (1999). Rapid ATM-dependent phosphorylation of MDM2 precedes p53 accumulation in response to DNA damage. Proceedings of the National Academy of Sciences of the United States of America, 96(26), 14973–14977. https://doi.org/10.1073/pnas.96.26.14973

Kim, S.-T., Xu, B., & Kastan, M. B. (2002). Involvement of the cohesin protein, Smc1, in Atm-dependent and independent responses to DNA damage. Genes & Development, 16(5), 560–570. https://doi.org/10.1101/gad.970602

Kitagawa, R., Bakkenist, C. J., McKinnon, P. J., & Kastan, M. B. (2004). Phosphorylation of SMC1 is a critical downstream event in the ATM-NBS1-BRCA1 pathway. Genes & Development, 18(12), 1423–1438. https://doi.org/10.1101/gad.1200304

Liao, H., Ji, F., Helleday, T., & Ying, S. (2018). Mechanisms for stalled replication fork stabilization: New targets for synthetic lethality strategies in cancer treatments. EMBO Reports, 19(9). https://doi.org/10.15252/embr.201846263

Lim, D. S., Kim, S. T., Xu, B., Maser, R. S., Lin, J., Petrini, J. H., & Kastan, M. B. (2000). ATM phosphorylates p95/nbs1 in an S-phase checkpoint pathway. Nature, 404(6778), 613–617. https://doi.org/10.1038/35007091

Luo, H., Li, Y., Mu, J.-J., Zhang, J., Tonaka, T., Hamamori, Y., Jung, S. Y., Wang, Y., & Qin, J. (2008). Regulation of intra-S phase checkpoint by ionizing radiation (IR)-dependent and IR-independent phosphorylation of SMC3. The Journal of Biological Chemistry, 283(28), 19176–19183. https://doi.org/10.1074/jbc.M802299200

Martin, S. L. (2006). The ORF1 Protein Encoded by LINE-1: Structure and Function During L1 Retrotransposition. Journal of Biomedicine and Biotechnology, 2006. https://doi.org/10.1155/JBB/2006/45621

Mathias, S. L., Scott, A. F., Kazazian, H. H., Boeke, J. D., & Gabriel, A. (1991). Reverse transcriptase encoded by a human transposable element. Science, 254(5039), 1808–1810. https://doi.org/10.1126/science.1722352

McDermott, J. E., Arshad, O. A., Petyuk, V. A., Fu, Y., Gritsenko, M. A., Clauss, T. R., Moore, R. J., Schepmoes, A. A., Zhao, R., Monroe, M. E., Schnaubelt, M., Tsai, C.-F., Payne, S. H., Huang, C., Wang, L.-B., Foltz, S., Wyczalkowski, M., Wu, Y., Song, E., … Rodland, K. D. (2020). Proteogenomic Characterization of Ovarian HGSC Implicates Mitotic Kinases, Replication Stress in Observed Chromosomal Instability. Cell Reports Medicine, 100004. https://doi.org/10.1016/j.xcrm.2020.100004

McKerrow, W., & Fenyö, D. (2020). L1EM: A tool for accurate locus specific LINE-1 RNA quantification. Bioinformatics (Oxford, England), 36(4), 1167–1173. https://doi.org/10.1093/bioinformatics/btz724

Mertins, P., Mani, D. R., Ruggles, K. V., Gillette, M. A., Clauser, K. R., Wang, P., Wang, X., Qiao, J. W., Cao, S., Petralia, F., Kawaler, E., Mundt, F., Krug, K., Tu, Z., Lei, J. T., Gatza, M. L., Wilkerson, M., Perou, C. M., Yellapantula, V., … NCI CPTAC. (2016). Proteogenomics connects somatic mutations to signalling in breast cancer. Nature, 534(7605), 55–62. https://doi.org/10.1038/nature18003

Miki, Y., Nishisho, I., Horii, A., Miyoshi, Y., Utsunomiya, J., Kinzler, K. W., Vogelstein, B., & Nakamura, Y. (1992). Disruption of the APC gene by a retrotransposal insertion of L1 sequence in a colon cancer. Cancer Research, 52(3), 643–645.

Minocherhomji, S., Ying, S., Bjerregaard, V. A., Bursomanno, S., Aleliunaite, A., Wu, W., Mankouri, H. W., Shen, H., Liu, Y., & Hickson, I. D. (2015). Replication stress activates DNA repair synthesis in mitosis. Nature, 528(7581), 286–290. https://doi.org/10.1038/nature16139

Mita, P., Sun, X., Fenyö, D., Kahler, D. J., Li, D., Agmon, N., Wudzinska, A., Keegan, S., Bader, J. S., Yun, C., & Boeke, J. D. (2020). BRCA1 and S phase DNA repair pathways restrict LINE-1 retrotransposition in human cells. Nature Structural & Molecular Biology, 27(2), 179–191. https://doi.org/10.1038/s41594-020-0374-z

Mita, P., Wudzinska, A., Sun, X., Andrade, J., Nayak, S., Kahler, D. J., Badri, S., LaCava, J., Ueberheide, B., Yun, C. Y., Fenyö, D., & Boeke, J. D. (2018, January 8). LINE-1 protein localization and functional dynamics during the cell cycle. ELife. https://doi.org/10.7554/eLife.30058

Morrison, C., Sonoda, E., Takao, N., Shinohara, A., Yamamoto, K., & Takeda, S. (2000). The controlling role of ATM in homologous recombinational repair of DNA damage. The EMBO Journal, 19(3), 463–471. https://doi.org/10.1093/emboj/19.3.463

Navarro, F. C., Hoops, J., Bellfy, L., Cerveira, E., Zhu, Q., Zhang, C., Lee, C., & Gerstein, M. B. (2019). TeXP: Deconvolving the effects of pervasive and autonomous transcription of transposable elements. BioRxiv, 648667. https://doi.org/10.1101/648667

Parker, J. S., Mullins, M., Cheang, M. C. U., Leung, S., Voduc, D., Vickery, T., Davies, S., Fauron, C., He, X., Hu, Z., Quackenbush, J. F., Stijleman, I. J., Palazzo, J., Marron, J. S., Nobel, A. B., Mardis, E., Nielsen, T. O., Ellis, M. J., Perou, C. M., & Bernard, P. S. (2009). Supervised risk predictor of breast cancer based on intrinsic subtypes. Journal of Clinical Oncology: Official Journal of the American Society of Clinical Oncology, 27(8), 1160–1167. https://doi.org/10.1200/JCO.2008.18.1370

Parrilla-Castellar, E. R., Arlander, S. J. H., & Karnitz, L. (2004). Dial 9-1-1 for DNA damage: The Rad9-Hus1-Rad1 (9-1-1) clamp complex. DNA Repair, 3(8–9), 1009–1014. https://doi.org/10.1016/j.dnarep.2004.03.032

Pellegrini, M., Celeste, A., Difilippantonio, S., Guo, R., Wang, W., Feigenbaum, L., & Nussenzweig, A. (2006). Autophosphorylation at serine 1987 is dispensable for murine Atm activation in vivo. Nature, 443(7108), 222–225. https://doi.org/10.1038/nature05112

Percharde, M., Lin, C.-J., Yin, Y., Guan, J., Peixoto, G. A., Bulut-Karslioglu, A., Biechele, S., Huang, B., Shen, X., & Ramalho-Santos, M. (2018). A LINE1-Nucleolin Partnership Regulates Early Development and ESC Identity. Cell, 174(2), 391–405.e19. https://doi.org/10.1016/j.cell.2018.05.043

Pizarro, J. G., & Cristofari, G. (2016). Post-Transcriptional Control of LINE-1 Retrotransposition by Cellular Host Factors in Somatic Cells. Frontiers in Cell and Developmental Biology, 4, 14. https://doi.org/10.3389/fcell.2016.00014

Rodić, N., Sharma, R., Sharma, R., Zampella, J., Dai, L., Taylor, M. S., Hruban, R. H., Iacobuzio-Donahue, C. A., Maitra, A., Torbenson, M. S., Goggins, M., Shih, I.-M., Duffield, A. S., Montgomery, E. A., Gabrielson, E., Netto, G. J., Lotan, T. L., De Marzo, A. M., Westra, W., … Burns, K. H. (2014). Long Interspersed Element-1 Protein Expression Is a Hallmark of Many Human Cancers. The American Journal of Pathology, 184(5), 1280–1286. https://doi.org/10.1016/j.ajpath.2014.01.007

Rodriguez-Martin, B., Alvarez, E. G., Baez-Ortega, A., Zamora, J., Supek, F., Demeulemeester, J., Santamarina, M., Ju, Y. S., Temes, J., Garcia-Souto, D., Detering, H., Li, Y., Rodriguez-Castro, J., Dueso-Barroso, A., Bruzos, A. L., Dentro, S. C., Blanco, M. G., Contino, G., Ardeljan, D., … PCAWG Consortium. (2020). Pan-cancer analysis of whole genomes identifies driver rearrangements promoted by LINE-1 retrotransposition. Nature Genetics, 52(3), 306–319. https://doi.org/10.1038/s41588-019-0562-0

Saldivar, J. C., Cortez, D., & Cimprich, K. A. (2017). The essential kinase ATR: Ensuring faithful duplication of a challenging genome. Nature Reviews. Molecular Cell Biology, 18(10), 622–636. https://doi.org/10.1038/nrm.2017.67

Scott, E. C., Gardner, E. J., Masood, A., Chuang, N. T., Vertino, P. M., & Devine, S. E. (2016). A hot L1 retrotransposon evades somatic repression and initiates human colorectal cancer. Genome Research, 26(6), 745–755. https://doi.org/10.1101/gr.201814.115

Smith, J., Tho, L. M., Xu, N., & Gillespie, D. A. (2010). The ATM-Chk2 and ATR-Chk1 pathways in DNA damage signaling and cancer. Advances in Cancer Research, 108, 73–112. https://doi.org/10.1016/B978-0-12-380888-2.00003-0

Taylor, M. S., Altukhov, I., Molloy, K. R., Mita, P., Jiang, H., Adney, E. M., Wudzinska, A., Badri, S., Ischenko, D., Eng, G., Burns, K. H., Fenyö, D., Chait, B. T., Alexeev, D., Rout, M. P., Boeke, J. D., & LaCava, J. (2018). Dissection of affinity captured LINE-1 macromolecular complexes. ELife, 7. https://doi.org/10.7554/eLife.30094

Taylor, M. S., LaCava, J., Mita, P., Molloy, K. R., Huang, C. R. L., Li, D., Adney, E. M., Jiang, H., Burns, K. H., Chait, B. T., Rout, M. P., Boeke, J. D., & Dai, L. (2013). Affinity proteomics reveals human host factors implicated in discrete stages of LINE-1 retrotransposition. Cell, 155(5), 1034–1048. https://doi.org/10.1016/j.cell.2013.10.021

Tubio, J. M. C., Li, Y., Ju, Y. S., Martincorena, I., Cooke, S. L., Tojo, M., Gundem, G., Pipinikas, C. P., Zamora, J., Raine, K., Menzies, A., Roman-Garcia, P., Fullam, A., Gerstung, M., Shlien, A., Tarpey, P. S., Papaemmanuil, E., Knappskog, S., Van Loo, P., … ICGC Prostate Cancer Group. (2014). Mobile DNA in cancer. Extensive transduction of nonrepetitive DNA mediated by L1 retrotransposition in cancer genomes. Science (New York, N.Y.), 345(6196), 1251343. https://doi.org/10.1126/science.1251343

Vasaikar, S., Huang, C., Wang, X., Petyuk, V. A., Savage, S. R., Wen, B., Dou, Y., Zhang, Y., Shi, Z., Arshad, O. A., Gritsenko, M. A., Zimmerman, L. J., McDermott, J. E., Clauss, T. R., Moore, R. J., Zhao, R., Monroe, M. E., Wang, Y.-T., Chambers, M. C., … Clinical Proteomic Tumor Analysis Consortium. (2019). Proteogenomic Analysis of Human Colon Cancer Reveals New Therapeutic Opportunities. Cell, 177(4), 1035–1049.e19. https://doi.org/10.1016/j.cell.2019.03.030

Vlastaridis, P., Kyriakidou, P., Chaliotis, A., Van de Peer, Y., Oliver, S. G., & Amoutzias, G. D. (2017). Estimating the total number of phosphoproteins and phosphorylation sites in eukaryotic proteomes. GigaScience, 6(2), 1–11. https://doi.org/10.1093/gigascience/giw015

Warkocki, Z., Krawczyk, P. S., Adamska, D., Bijata, K., Garcia-Perez, J. L., & Dziembowski, A. (2018). Uridylation by TUT4/7 Restricts Retrotransposition of Human LINE-1s. Cell, 174(6), 1537–1548.e29. https://doi.org/10.1016/j.cell.2018.07.022

Wei, W., Gilbert, N., Ooi, S. L., Lawler, J. F., Ostertag, E. M., Kazazian, H. H., Boeke, J. D., & Moran, J. V. (2001). Human L1 Retrotransposition: Cis Preference versus trans Complementation. Molecular and Cellular Biology, 21(4), 1429–1439. https://doi.org/10.1128/MCB.21.4.1429-1439.2001

Wiredja, D. D., Koyutürk, M., & Chance, M. R. (2017). The KSEA App: A web-based tool for kinase activity inference from quantitative phosphoproteomics. Bioinformatics (Oxford, England), 33(21), 3489–3491. https://doi.org/10.1093/bioinformatics/btx415

Wylie, A., Jones, A. E., D’Brot, A., Lu, W.-J., Kurtz, P., Moran, J. V., Rakheja, D., Chen, K. S., Hammer, R. E., Comerford, S. A., Amatruda, J. F., & Abrams, J. M. (2016). P53 genes function to restrain mobile elements. Genes & Development, 30(1), 64–77. https://doi.org/10.1101/gad.266098.115

Yang, F., & Wang, P. J. (2016). Multiple LINEs of retrotransposon silencing mechanisms in the mammalian germline. Seminars in Cell & Developmental Biology, 59, 118–125. https://doi.org/10.1016/j.semcdb.2016.03.001

Yazdi, P. T., Wang, Y., Zhao, S., Patel, N., Lee, E. Y.-H. P., & Qin, J. (2002). SMC1 is a downstream effector in the ATM/NBS1 branch of the human S-phase checkpoint. Genes & Development, 16(5), 571–582. https://doi.org/10.1101/gad.970702

Yoshihara, K., Shahmoradgoli, M., Martínez, E., Vegesna, R., Kim, H., Torres-Garcia, W., Treviño, V., Shen, H., Laird, P. W., Levine, D. A., Carter, S. L., Getz, G., Stemke-Hale, K., Mills, G. B., & Verhaak, R. G. W. (2013). Inferring tumour purity and stromal and immune cell admixture from expression data. Nature Communications, 4, 2612. https://doi.org/10.1038/ncomms3612

Yu, Q., Carbone, C. J., Katlinskaya, Y. V., Zheng, H., Zheng, K., Luo, M., Wang, P. J., Greenberg, R. A., & Fuchs, S. Y. (2015). Type I interferon controls propagation of long interspersed element-1. The Journal of Biological Chemistry, 290(16), 10191–10199. https://doi.org/10.1074/jbc.M114.612374

Zhang, H., Liu, T., Zhang, Z., Payne, S. H., Zhang, B., McDermott, J. E., Zhou, J.-Y., Petyuk, V. A., Chen, L., Ray, D., Sun, S., Yang, F., Chen, L., Wang, J., Shah, P., Cha, S. W., Aiyetan, P., Woo, S., Tian, Y., … Townsend, R. R. (2016). Integrated Proteogenomic Characterization of Human High-Grade Serous Ovarian Cancer. Cell, 166(3), 755–765. https://doi.org/10.1016/j.cell.2016.05.069

